# Vascular Patterning Shapes Intramembranous Ossification via HIF1α-VEGF Axis

**DOI:** 10.1101/2025.09.13.676037

**Authors:** Soma Dash, Jonathan R. Rettig, Madelaine Gogol, Paul A. Trainor

**Affiliations:** Department of Biological Sciences, University at Albany, SUNY Albany; RNA Institute, University at Albany, SUNY Albany; Stowers Institute for Medical Research; University of Kansas, Medical Center

**Author notes:** Corresponding author: Soma Dash. Data availability: http://www.stowers.org/research/publications/LIBPB-2590 https://doi.org/10.7910/DVN/I0QP2C https://github.com/mmarchin/cbio.sdash.101.

**Keywords:** Mediator complex, Vascular development, Craniofacial Development, Spatial transcriptomics, HIF1α, VEGF signaling

## Abstract

Organs and tissues develop in close association with vasculature which transports blood and nutrients and helps to remove waste. The vasculature is composed primarily of endothelial cells, which provide structure, form barriers, and are a source of developmental signals. We recently found that the Mediator, a multiprotein complex, which regulates transcription, was essential for proper vascular development. Here, we investigated the specific role of the Mediator tail subunit Med23 in endothelial cells. Endothelial cell-specific knockout of *Med23* in mouse embryos using *Tek-Cre* results in vascular anomalies, including edema, hemorrhage, and mispatterned vasculature, alongside craniofacial defects such as micrognathia and cleft palate. Spatial transcriptomics revealed downregulated expression of key vascular and osteogenic genes in *Med23* mutants, including *Vegfr1* and *Col1a1*, with altered signaling dynamics between endothelial and osteoblast populations. Elevated HIF1α expression and reduced VEGF signaling were observed in *Med23* mutants, suggesting a hypoxia-driven suppression of neural crest cell derived osteoblast maturation. Consistent with this model, pharmacological inhibition of HIF1α, combined with VEGFA supplementation, rescued craniofacial ossification and extended embryonic viability. These findings reveal a critical role for Med23 in coordinating vascular patterning and intramembranous ossification and highlight distinct hypoxic and angiogenic requirements in craniofacial dermal bone versus axial and appendicular endochondral bone development. Thus, the cranial vasculature and more specifically endothelial cells, play an instructive role in neural crest cell and osteogenic differentiation during cranioskeletal development.

## Introduction

Bone development and repair are intimately linked to vascularization, with endothelial cells (ECs) providing angiocrine signals that support both osteogenesis and hematopoiesis (Sivan et al. 2019; Zhu et al. 2020). Although the vascular dynamics of endochondral ossification are well characterized, the role of vasculature in intramembranous ossification remains poorly understood (Percival and Richtsmeier 2013). Notably, both ossification modes require avascular conditions at the initiation of primary ossification centers, followed by angiogenesis to sustain bone expansion (Thompson et al. 1989). This vascular influx not only supplies metabolic support but also ECs that interact with osteoblast precursors to regulate their differentiation and survival. ECs in the vasculature, depending on their proximity to the osteoblasts, have been shown to respond to pre-osteoblast cells, which then through various signaling pathways regulate the differentiation and survival of osteoblast cells (Collin-Osdoby 1994; Brandi and Collin-Osdoby 2006).

Osteoblasts and progenitors reciprocally influence EC behavior by secreting pro-angiogenic factors, including VEGF ligands. Hypoxia-induced VEGF signaling, mediated by HIF1α stabilization, is a key driver of both angiogenesis and osteogenesis (Zelzer et al. 2002; Zelzer et al. 2004). HIF1α, a master regulator of the cellular response to hypoxia, promotes VEGF expression and enhances osteoblast differentiation in long bones under low oxygen conditions (Wang et al. 2007). During development and regeneration through the endochondral ossification process, HIF1α is essential for coupling angiogenesis with osteogenesis (Shao et al. 2022).

VEGF signaling plays a role in both angiogenesis and osteogenesis and its levels must be tightly regulated to ensure proper development (Grosso et al. 2023). Among the five VEGF ligands, Vegfa is the most abundant and functionally versatile, signaling primarily through Vegfr2 (Flk1, Kdr), while Vegfr1 (Flt1) acts as a decoy receptor reducing angiogenesis (Waltenberger et al. 1994; Meyer et al. 2006). Regulation of VEGF signaling is tissue-specific and complex, involving transcriptional control by Runx2 and potential modulation by FGF and BMP pathways (Zelzer et al. 2001).

Recent studies have highlighted the Mediator complex—a central integrator of transcriptional regulation—as a key player in craniofacial and vascular development (Hashimoto et al. 2011; Liu et al. 2016; Dash et al. 2020; Dash et al. 2021; Yang et al. 2022). The subunit Med23, in particular, has emerged as a critical regulator of EC function and neural crest cell NCC) during craniofacial morphogenesis (Dash et al. 2020; Dash et al 2021). Loss of Med23 in NCC results in craniofacial anomalies similar to Pierre-Robin sequence, including cleft palate and micrognathia, likely due to perturbed modulation of WNT and Sox9 signaling (Dash et al. 2021). These findings underscore Med23’s role as a molecular bridge linking transcriptional control to morphogenetic and angiogenic processes during embryogenesis.

In this study, we describe an EC-specific Med23 knockout mouse that survives to embryonic day (E)16.5 and exhibits both vascular and craniofacial anomalies. Although, Med23 deletion in ECs has been previously shown to cause hemorrhage and reduced angiogenesis, (Yang et al. 2022) differences in allele design allow our mutants to survive longer, enabling analysis of cranioskeletal phenotypes in addition to vascular defects. Together with edema and hemorrhage arising from disruption of the HIF1α–VEGF signaling axis, we observe impaired craniofacial osteogenesis, evident by reduced expression of key osteogenic markers. Consistent with these observations, pharmacological inhibition of HIF1α, combined with VEGF pathway activation, rescues the craniofacial ossification phenotype. Together, these findings support a model in which Med23 coordinates vascular and craniofacial development by regulating the spatial and temporal dynamics of ossification through vascular-mediated hypoxic signaling.

## Results

### Tissue specific deletion of Med23 from EC results in vascular anomalies

In a forward genetic screen aimed at identifying novel regulators of craniofacial development (Sandell et al. 2011), we discovered a mouse mutant, *snouty*, which exhibited craniofacial and neurovascular anomalies but was early embryonically lethal (Dash et al. 2020; Dash et al. 2021). We subsequently mapped the *snouty* mutation to a single nucleotide change in a ubiquitously expressed gene, *Med23* (Dash et al. 2020), which encodes a subunit of the global transcription co-factor complex, Mediator. To overcome the early embryonic lethality of *Med23*^*sn/sn*^ mutants and investigate the role of Med23 in later vascular development and maturation, we conditionally deleted *Med23* specifically in ECs with *Tek-Cre* (*Tie2-Cre*) transgenic mice.

*Med23*^*fx/fx*^*;Tek-Cre* mutant embryos (hereby referred to as *Med23*^*ECKO/ECKO*^) survived until late gestation at E16.5 but presented with edema at E14.5 (Fig. 1A, A’, E, E’) that progressively worsened (Fig. 1B, B’). To investigate the developmental origin of these vascular anomalies, we performed lineage tracing of EC precursors using *Rosa-eYFP* reporter mice.

**Fig. 1:**
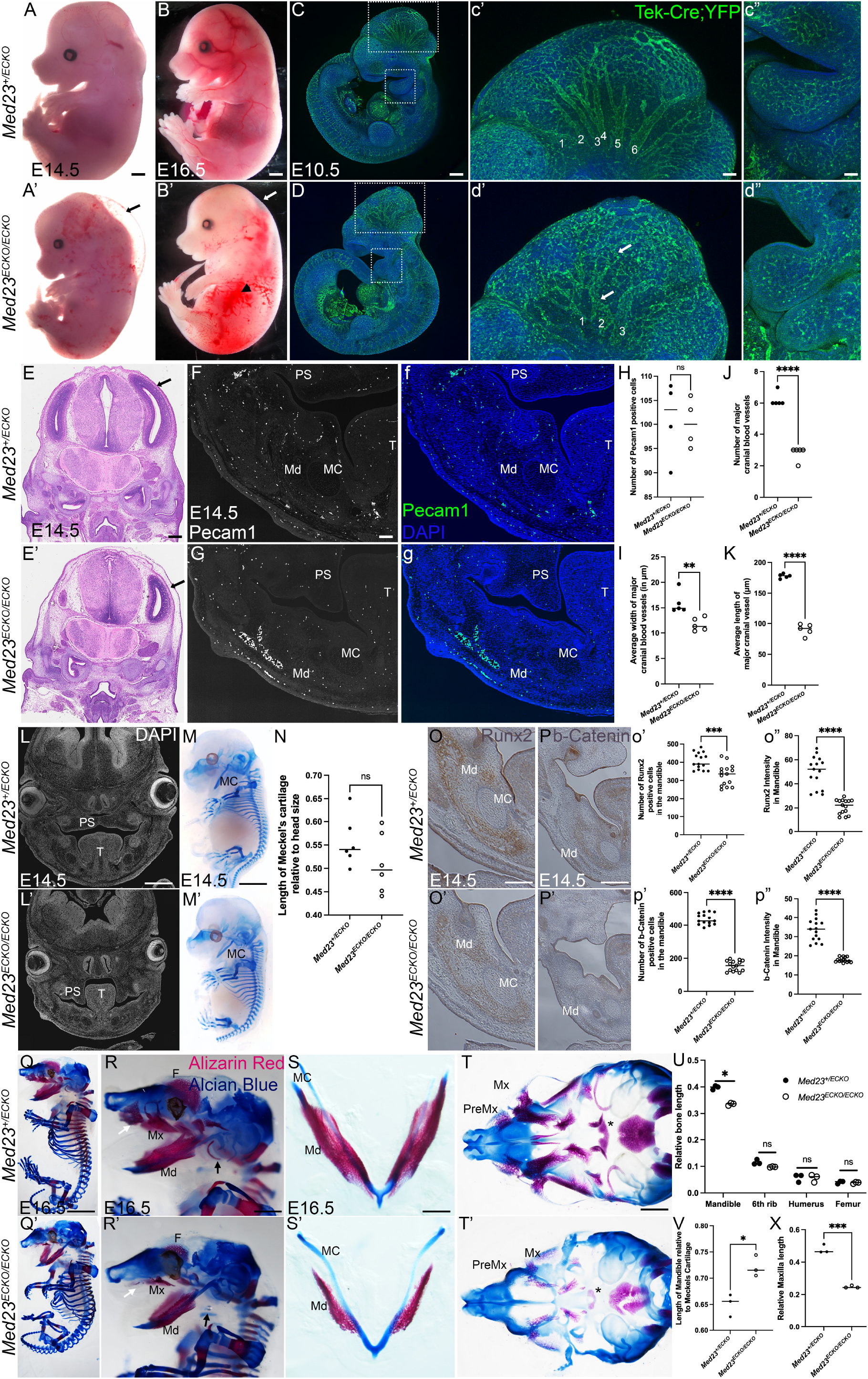
Vascular and craniofacial anomalies in *Med23*^*ECKO/ECKO*^ mutants. Bright field images of *Med23*^*+/ECKO*^ and *Med23*^*ECKO/ECKO*^ at E14.5 (A,A’) and E16.5 (B,B’) indicate that mutants exhibit vascular defects such as edema (indicated by arrows) and hemorrhage (arrowhead in B’) (n=10). Scale bar in A,A = 500 μm, B’B’ = 600 μm, Lineage tracing using a Rosa-eYFP transgenic line followed by staining with GFP antibody indicates that the vascular patterning was disrupted in *Med23*^*ECKO/ECKO*^ compared to *Med23*^*+/ECKO*^ (n=5) (C,D). Scale bar = 150 μm. A high magnification image of the brain region and pharyngeal arches are shown in c’ and d’ and c” and d”, respectively. Arrows indicate the disrupted vasculature in *Med23*^*ECKO/ECKO*^ in d’. Scale bar in c’,d’ = 50 μm, c”,d” = 20 μm. Histological analysis of E14.5 embryos indicate that the blister observed in B is a fluid filled space, empty of cells (indicated by black arrow) (E, E’). Scale bar = 200 μm. Pecam1 immunostaining in white (F, G) and green (f, g) on E14.5 sections indicates that the number of Pecam1 positive cells in the lower jaw is similar in *Med23*^*+/ECKO*^ and *Med23*^*ECKO/ECKO*^ (n=4). Scale bar = 140 μm. Quantification of Pecam1 positive cells in the lower jaw is shown in K. Quantification of the width, number and length of major cranial blood vessels at E10.5 (numbered in E’ and F’) are shown in L, M and N, respectively. (L,L’) Sectioned and DAPI stained E14.5 embryos indicate that while the palatal shelf in the *Med23*^*+/ECKO*^ control embryos are fused in a horizontal position above the tongue, *Med23*^*ECKO/ECKO*^ mutants exhibit cleft palate with palatal shelves in a vertical position flanking the tongue (n=10). Scale bar = 150 μm. (M,M’) Alcian blue staining reveals no substantial difference in cartilage morphology between *Med23*^*+/ECKO*^ and *Med23*^*ECKO/ECKO*^ at E14.5 (n=5). Scale bar = 200 μm. (N) Quantification of the normalized size of the Meckel’s cartilage compared to the size of the head (tip of the snout to hindbrain). Osteoblast progenitors labeled by Runx2 (O,O’,o’,o”) and b-Catenin (P,P’,p’,p”) indicate significantly reduced expression of both Runx2 and b-Catenin in the mandible (n = 5). Scale bar = 75 μm. (Q,Q’) Alcian blue and alizarin red staining of E16.5 *Med23*^*+/ECKO*^ and *Med23*^*ECKO/ECKO*^ embryos indicates the presence of craniofacial osteogenesis defects (n=3). Scale bar = 300 μm. High magnification images of the craniofacial region displayed in R and R’ show that the frontal bone, premaxilla (indicated by white arrowheads), maxilla and mandible are drastically reduced in *Med23*^*ECKO/ECKO*^ embryos, while the tympanic bone is missing in the *Med23*^*ECKO/ECKO*^ embryos (indicated by black arrowheads). Scale bar = 400 μm. (S,S’) Reduced alizarin red staining in dissected lower jaw indicates that the mandible ossification is reduced in *Med23*^*ECKO/ECKO*^ embryos compared to *Med23*^*+ECKO*^. Scale bar = 75 μm. (T,T’) Reduced alizarin red staining in dissected upper jaw indicates that the ossification in premaxilla, maxilla and palatine bone is reduced in *Med23*^*ECKO/ECKO*^ embryos compared to *Med23*^*+ECKO*^. Scale bar = 200 μm. (U) Quantification of relative mineralized bone length compared to the body length (crown to rump) indicates that other than the craniofacial region, no bones are altered significantly in size. Mandible length was normalized to the size of the head (nose to hind brain). (V) Quantification of the mandible length relative to Meckel’s cartilage length. (X) Quantification of the normalized size of the maxilla compared to the head size. Modified t-test was used for statistical analyses. Abbr. PS - Palatal Shelf, T – Tongue, MC - Meckel’s Cartilage, Md – Mandible, Mx - Maxilla, PreMx – Premaxilla, F-Frontal bone.

While the vascular network was present in E10.5 *Med23*^*ECKO/ECKO*^ mutants, its patterning was notably altered particularly in the forebrain, midbrain and pharyngeal arches compared to *Med23*^*+/ECKO*^ control embryos (Fig. 1C-d”). In contrast to the reiterated tree branch-like pattern of large dorsoventrally oriented vessels connected to a well-organized polygonal network of smaller vessels in controls, the major vessels of the midbrain associated vascular plexus in *Med23*^*ECKO/ECKO*^ mutants were less directional, disorganized, fewer in number, and had reduced width and length (Fig. 1I-K), which was collectively indicative of perturbed vascular remodeling. In the mandible, the number of Pecam1-positive ECs appeared comparable between *Med23*^*+/ECKO*^ control and *Med23*^*ECKO/ECKO*^ mutant embryos (Fig. 1F-H), however their pattern was disrupted.

### Tissue specific deletion of Med23 from ECs results in craniofacial anomalies

Closer examination of *Med23*^*ECKO/ECKO*^ mutants revealed craniofacial anomalies, including micrognathia (Fig. 1A’, B’) and cleft palate (Fig. 1L, L’, Appendix Fig. 1). To further characterize these defects, we performed Alizarin red and Alcian blue skeletal staining of bone and cartilage respectively. At E14.5, craniofacial and axial cartilage development appeared normal in *Med23*^*ECKO/ECKO*^ mutants compared to *Med23*^*+/ECKO*^ controls, including the length of Meckel’s cartilage suggesting that Med23 ECs is not essential in ECs for craniofacial chondrogenesis (Fig. 1M, M’). In contrast, Runx2, a master regulator of osteogenic differentiation (Otto et al. 1997), and β-Catenin, a Wnt signaling effector molecule necessary for craniofacial bone formation (Brault et al. 2001), were both markedly reduced in the mandibular region of *Med23*^*ECKO/ECKO*^ mutants (Fig. 1O-p”), consistent with the decrease in Alizarin red staining.

By E16.5, while cartilage differentiation and growth continued to be unaffected, craniofacial osteogenesis was markedly reduced in *Med23*^*ECKO/ECKO*^ mutant embryos, resulting in hypoplasia of the frontal, premaxillary, maxillary, tympanic, palatine and mandibular bones (Fig. 1Q-R’). These phenotypes are fully penetrant; however, the severity of the vascular defects varies among mutants, with approximately 25% exhibiting early physiological decline by E16.5.

To assess the regional specificity of skeletal differentiation, we compared osteogenesis in the ribs and long bones to that in the craniofacial region. Notably, intramembranous ossification including that of the mandible and maxilla was markedly reduced, while skeletal elements formed via endochondral ossification were unaffected (Fig. 1S-X). These findings suggest that vascular mispatterning resulting from *Med23* deletion in ECs selectively impairs craniofacial intramenbranous osteogenesis, likely through disrupted signaling between the vasculature and osteoprogenitor cells. We use the mandible as a representative model system of intramembranous ossification for all subsequent analysis.

### NCC migration is minimally affected in the *Med23*^*ECKO/ECKO*^ mutants

Since most craniofacial bones are derived from NCC, we examined NCC migration at E10.5 using Sox9 and Sox10, which are also master regulators of cartilage and bone, and neuro-glial progenitors respectively (Kuhlbrodt et al. 1998; Bi et al. 1999). Expression of each gene was comparable between *Med23*^*+/ECKO*^ control and *Med23*^*ECKO/ECKO*^ mutant embryos with Sox9 in the pharyngeal arches and Sox10 in the trigeminal region, indicating that NCC induction and migration were not disrupted (Appendix Fig. 2A-F). To assess overall cranial nerve development, we performed β-Tubulin III staining and observed intact and properly patterned cranial nerves and dorsal root ganglia in both controls and mutants (Appendix Fig. 2G,H), demonstrating that NCC differentiation into neurons was unaffected by the alterations in the vasculature. Finally, we quantified the volume of the mandibular arch and observed that at E10.5, its size is comparable between *Med23*^*+/ECKO*^ control and *Med23*^*ECKO/ECKO*^ mutant embryos (Appendix Figure 2I).

These findings suggest that although NCCs successfully migrate to the mandibular arch, their differentiation into osteochondroprogenitors may be non-cell autonomously compromised by the absence of Med23 in ECs, which implicates vascular signaling in the regulation of intramembranous ossification during cranioskeletal development.

### Proliferation and Apoptosis Anomalies in *Med23*^*ECKO/ECKO*^ Mutants

Considering craniofacial osteogenesis is markedly reduced in *Med23*^*ECKO/ECKO*^ mutants, we hypothesized that the osteochondroprogenitor population was reduced due to decreased proliferation or increased apoptosis. To assess cellular proliferation and apoptosis in *Med23*^*ECKO/ECKO*^ mutants, we performed immunostaining with antibodies against Phospho-Histone H3 (pHH3) and Cleaved Caspase-3 (CC3), respectively on coronal sections of E12.5 and E14.5 embryos. We observed no changes in cell proliferation but detected a slight increase in apoptosis of vasculature associated ECs in sections of E14.5 *Med23*^*ECKO/ECKO*^ mutants compared to *Med23*^*+/ECKO*^ controls, but not in the mandibular mesenchyme at either E12.5 or E14.5 (Appendix Fig.3).

### Spatial transcriptomics reveals transcriptomic changes in the osteochondroprogenitor cells

To investigate the interactions between osteochondroprogenitor cells and the vasculature, we performed spatial transcriptomics using the CosMx platform (NanoString), which allowed us to profile 990 distinct transcripts across diverse cell types. We analyzed two coronal tissue sections each from two *Med23*^*+/ECKO*^ control and two *Med23*^*ECKO/ECKO*^ mutant embryos. Data were processed using the Seurat package in R and visualized via UMAP clustering to identify cell types based on gene expression, which were then mapped back to histological landmarks (Fig. 2A-C). This analysis revealed two distinct vascular cell populations, classified as ECs and pericytes and two distinct osteoblast populations, classified as immature and mature osteoblasts.

**Fig. 2:**
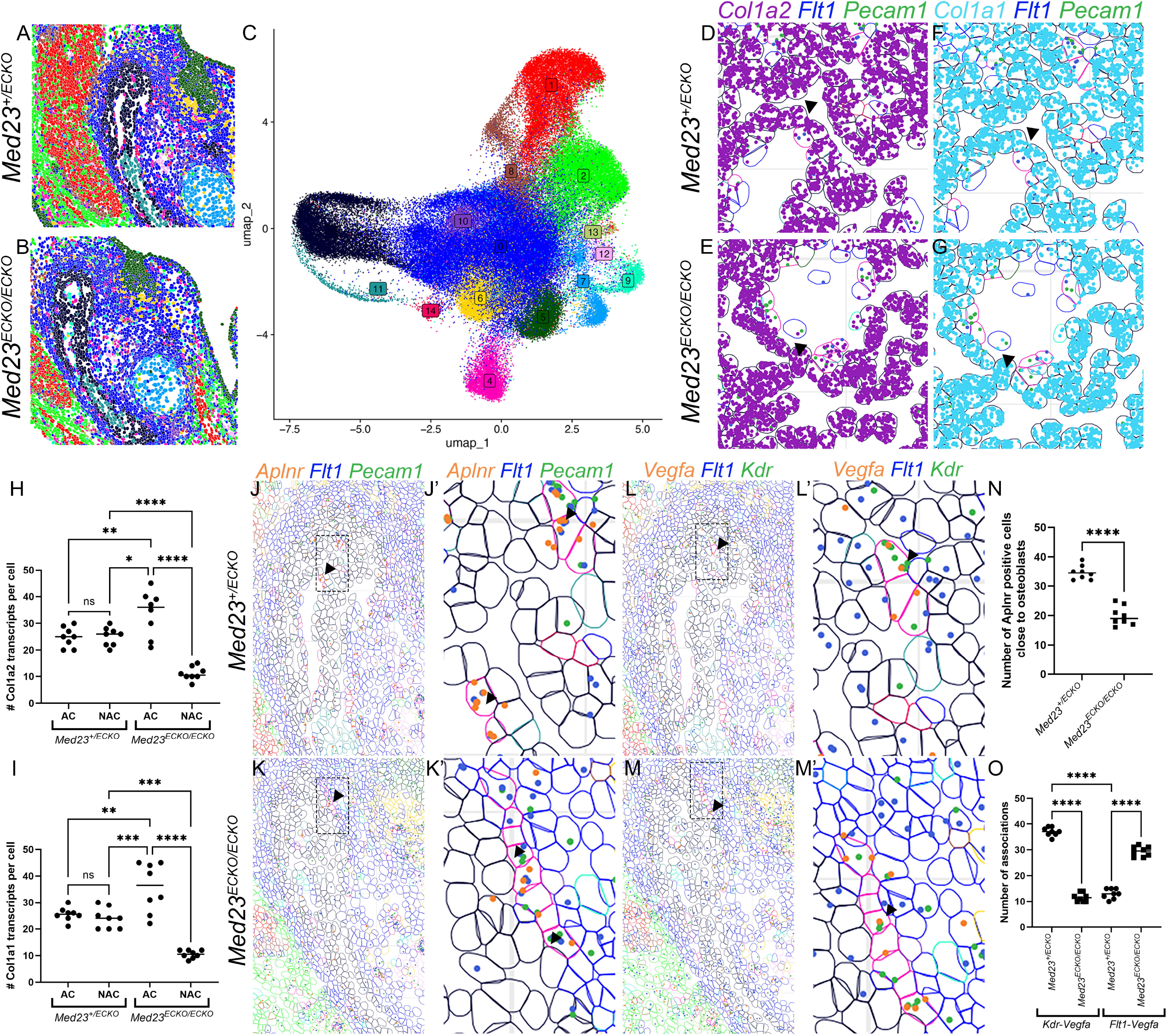
Spatial Transcriptomics indicates disruption of osteogenesis and angiogenesis genes. (A-C) Spatial transcriptomics identifies tissue based on gene expression patterns of 990 genes that comprise a whole-body panel on the NanoString CosMx platform. Uniform Manifold Approximation and Projection (UMAP) clustering and a feature plot of the major tissue types in the mandibular region led to the identification of 14 cell populations: Mesenchyme 1 (0, blue), Masseter Muscle (1, red), Mesenchyme 2 (2, light green), Immature Osteoblasts (3, black), Endothelial Cells (4, dark pink), Tooth (5, dark green), Dental Mesenchyme (6, yellow), Meckel’s Cartilage (7, light blue), Myogenic Progenitors (8, brown), Glia (9, teal), Neuron 1 (10, purple), Mature Osteoblasts (11, dark teal), Neuron 2 (12, light pink), Epithelial cells (13, olive green), and Pericytes (14, dark red). Spatial transcriptomics-based expression of *Col1a2* (purple, D-E) and *Col1a1* (cyan, F-G) in the adjoining (AC) osteoblasts (black outlined) compared to non-adjoining (NAC) endothelial cells (pink outlined) in *Med23*^*+/ECKO*^ controls and *Med23*^*ECKO/ECKO*^ mutants. Quantification of *Col1a2* transcripts (H) and *Col1a1* transcripts (I) were statistically analyzed using ANOVA. (J-K’,N) Spatial transcriptomics-based expression of *Aplnr* (orange) in conjunction with *Flt1* (*Vegfr1*, blue) and *Pecam1* (green) around the mandible. (L-M’) *Vegfa* (orange), *Flt1* (blue) and *Kdr* (*Vegfr2*, green) expression was analyzed and used to generate figures in R using Seurat. (O) Quantification of the number of *Vegfa* positive cells in proximity to *Vegfr2* positive cells compared to *Vegfa* positive cells in proximity to *Vegfr1* positive cells in *Med23*^*+/ECKO*^ controls and *Med23*^*ECKO/ECKO*^ mutants. Statistical analysis was performed using ANOVA.

Differential gene expression analysis (padj < 0.05, ≥20% expression in either group, fold change >1.5) revealed three significantly altered genes in vascular cells: *Gnas, Aplnr*, and *Flt1* (Appendix Table 1). *Gnas* encodes the stimulatory G protein alpha subunit, which when mutated causes pseudohypoparathyroidism leading to vascular calcification (Klenke et al. 2020) and defects in the cranial base, maxilla and mandible (Le Norcy et al. 2020). *Aplnr*, which encodes the Apelin receptor, is essential for vasculogenesis and angiogenesis (Helker et al. 2020). *Aplnr* expression was downregulated in *Med23*^*ECKO/ECKO*^ mutants, likely contributing to the vascular mispatterning (Fig. 2J-K’,N). *Flt1* (VEGFR1), a key modulator of angiogenesis through competion with VEGFR2 for VEGFA, was upregulated in *Med23*^*ECKO/ECKO*^ mutants, while *Vegfa* transcript levels remained unchanged (Fig. 2L-M’). However, more EC in the mutants express *Vegfa* consistent with disrupted vascular organization and potentially hypoxia.

With respect to osteoblasts, key matrix protein encoding genes which are also required for osteogenic differentiation including *Col1a1, Col1a2*, and *Sparc* (Rodriguez Celin et al. 1993; Mendoza-Londono et al. 2015), were downregulated together with β-Catenin in mature osteoblasts (Appendix Table 1). Additionally, downregulation of *Thy1* and *Gja1* expression, which encode cell adhesion proteins that are important for craniofacial mesenchymal differentiation (Paznekas et al. 2003; An et al. 2018; Andrade Azevedo de Vasconcelos et al. 2025), further indicate osteoblast maturation in the mutants is impaired—consistent with the skeletal phenotypes evident in *Med23*^*ECKO/ECKO*^ mutants.

Spatial analysis revealed a striking pattern in *Med23*^*+/ECKO*^ control embryos, in which immature osteoblasts adjacent to and distant from vasculature expressed similar levels of osteogenic genes. In contrast, in *Med23*^*ECKO/ECKO*^ mutants, osteoblasts near vasculature exhibited elevated expression of *Col1a1* and *Col1a2*, while distant cells exhibited reduced expression of *Col1a1* and *Col1a2* (Fig. 2D-I). This is suggestive of enhanced osteogenesis in the vicinity of the vasculature in *Med23*^*ECKO/ECKO*^ mutants.

Interestingly, CellChat analysis of the spatial transcriptomics data revealed altered Vegf signaling dynamics (Fig. 2O). Vegfa–Vegfr2 interactions were diminished in *Med23*^*ECKO/ECKO*^ mutants, while Vegfa–Vegfr1 interactions were enhanced compared to *Med23*^*+/ECKO*^ controls, supporting a shift in the balance of VEGF signaling that may underlie both vascular and osteogenic defects.

### Hypoxia is increased in *Med23*^*ECKO/ECKO*^ mutants

VEGF signaling must be downregulated in long bones to promote osteoblast maturation and repair, a process mediated by hypoxia induced HIF1α signaling. We therefore investigated HIF1α signaling in *Med23*^*ECKO/ECKO*^ mutants and observed a significant upregulation of HIF1α expression, particularly around the dentary bone, which is suggestive of elevated hypoxia in *Med23*^*ECKO/ECKO*^ mutants compared to *Med23*^*+/ECKO*^ controls (Fig. 3A-E).

**Fig 3:**
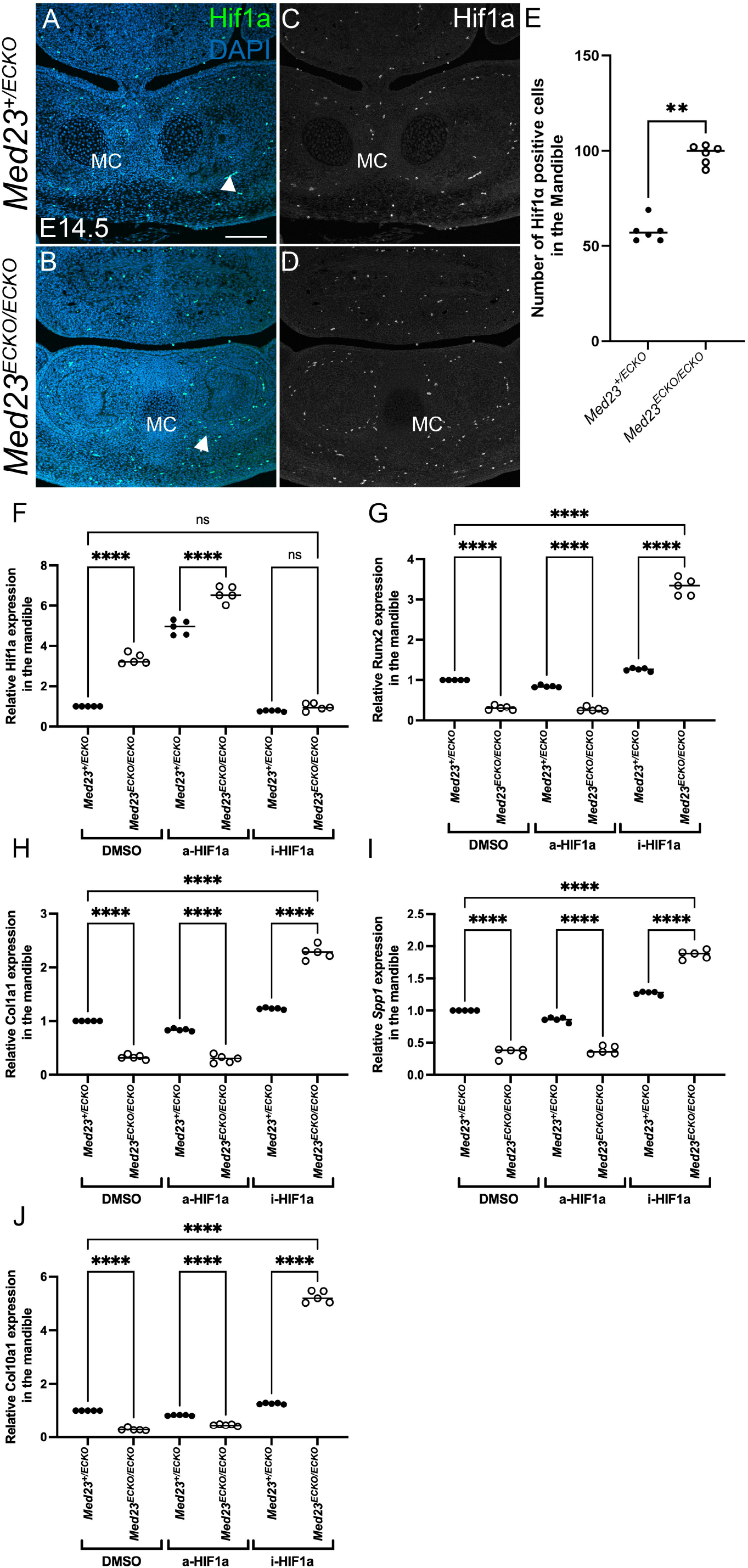
Hypoxia gene is upregulated in *Med23*^*ECKO/ECKO*^ mutants. (A-D) Coronal sections of E14.5 *Med23*^*+/ECKO*^ control and *Med23*^*ECKO/ECKO*^ mutant embryos were immunostained for HIF1α. Scale bar: 140 μm. (E) Quantification of HIF1α-positive cells in the mandible. Statistical analysis was performed using a non-parametric t-test; n = 6. qPCR with isolated RNA from *Med23*^*+/ECKO*^ controls and *Med23*^*ECKO/ECKO*^ embryos treated with either activator or inhibitor of HIF1α compared to DMSO treated embryos for Hif1a (F), Runx2 (G), Col1a1 (H), Spp1 (I) and Col10a1 (J).

To analyze the effect of HIF1α upregulation in *Med23*^*ECKO/ECKO*^ mutants, we pharmacologically treated pregnant dams with a HIF1α activator (DMOG, a-HIF1a) and inhibitor (FM19G11, i-HIF1a) and then collected the embryos for qPCR analysis. When treated with a-HIF1a, Hif1α was increased in control and mutant embryos as expected. However, a-HIF1a *Med23*^*ECKO/ECKO*^ mutant embryos did not exhibit altered *Runx2, Col1a1, Spp1* and *Col10a1* in comparison to *Med23*^*+/ECKO*^ control vehicle treated embryos, suggesting no effect of HIF1a activation on the state of osteogenic differentiation. In contrast, when treated with the i-HIF1a, both *Med23*^*+/ECKO*^ control and *Med23*^*ECKO/ECKO*^ mutant explants exhibited increased expression of osteogenic genes in comparison to *Med23*^*+/ECKO*^ and *Med23*^*ECKO/ECKO*^ mutant treated with DMSO, consistent with enhanced osteogenesis (Fig. 3F-J) .

### VEGF-HIF1α modulation rescues osteogenesis in *Med23*^*ECKO/ECKO*^ mutants

Our data collectively suggested that elevated HIF1α and altered VEGF signaling perturbed craniofacial bone formation in *Med23*^*ECKO/ECKO*^ mutants. To test this idea, we performed pharmacological inhibition and activation of VEGF signaling in combination with inhibition of HIF1α signaling to investigate the mechanistic effects of VEGF-HIF1α signaling on intramembranous ossification. As expected, VEGF signaling inhibition using VEGF inhibitor ZM323881 disrupted osteogenesis in *Med23*^*+/ECKO*^ control embryos compared to DMSO carrier controls. However, *Med23*^*ECKO/ECKO*^ mutant embryos exhibited minimal additional disruption, likely becuase VEGF signaling is already significantly reduced in *Med23*^*ECKO/ECKO*^ mutants such that further VEGF signaling suppression does not exacerbate the phenotype (Fig. 4A-H).

**Fig. 4:**
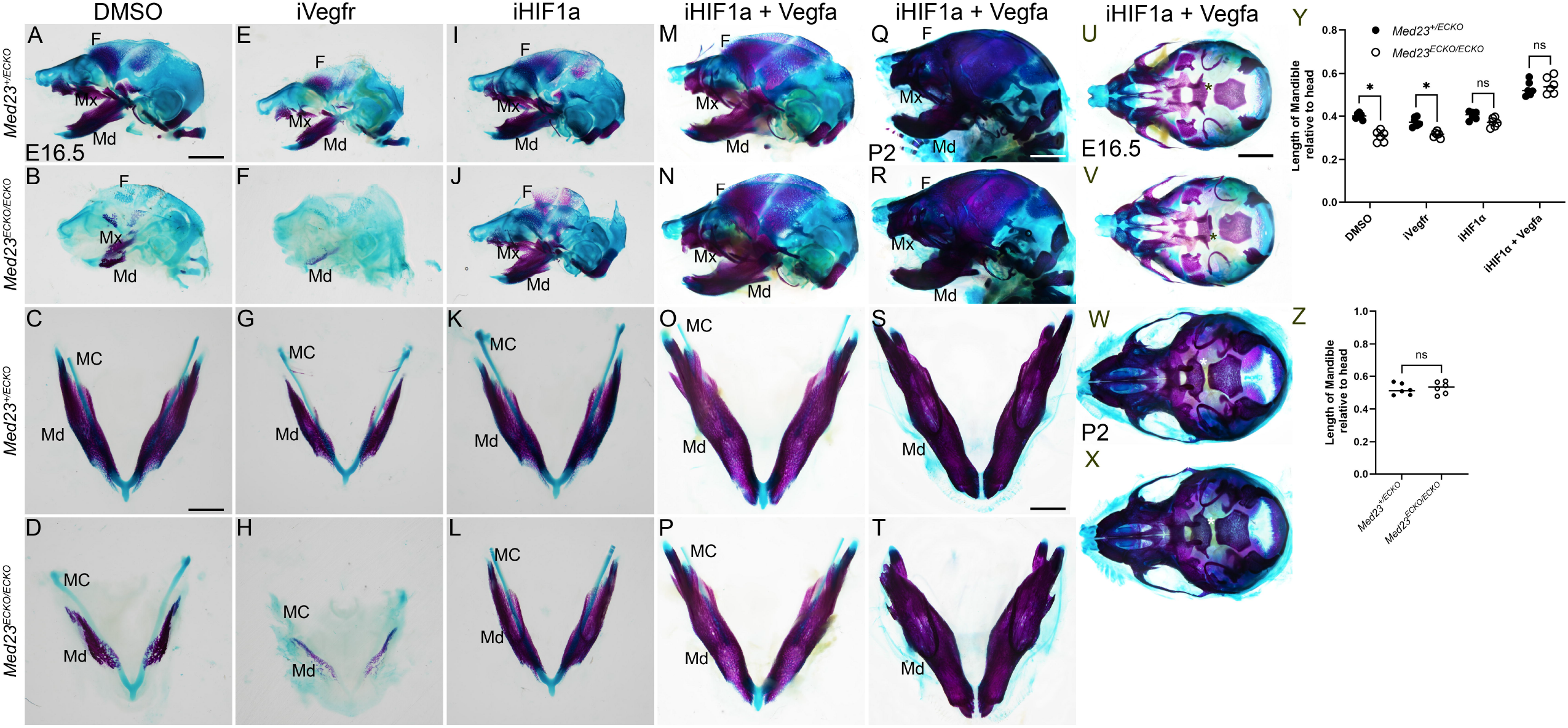
Osteogenesis rescue in *Med23*^*ECKO/ECKO*^ mutants with Vegf-HIF1a modulation. Alcian blue and alizarin red staining of *Med23*^*+/ECKO*^ controls and *Med23*^*ECKO/ECKO*^ mutants treated with DMSO (A-D), Vegfr inbibitor (iVegfr, E-H), Hif1a inhibitor (iHIF1a, I-L) and Hif1a inhibitor combined with Vegfa (iHIF1a + Vegfa, M-P, U-V) at E16.5 reveals partial rescue of craniofacial osteogenesis using iHIF1a and complete rescue with iHIF1a + Vegfa. (Q-T,W-X) *Med23*^*ECKO/ECKO*^ mutant embryos treated with iHIF1a + Vegfa survive until P2 unlike DMSO treated mutants. At P2, the mandible size along with other craniofacial bones are comparable between *Med23*^*+/ECKO*^ controls and *Med23*^*ECKO/ECKO*^ mutants indicating complete rescue of the mutants. (Y) Quantification of length of the mandible relative to the length of the head (n=6) of E16.5 embryos treated with DMSO, iVegfr, iHIF1a and iHIF1a + Vegfa. Mann-Whitney U test was performed for statistical analysis. (Z) Quantification of the length of the mandible relative to the length of the head (n=6) treated with iHIF1a + Vegfa of both *Med23*^*+/ECKO*^ controls and *Med23*^*ECKO/ECKO*^ mutants at P2. Scale bar for A,B, E, F, I, J, M, N is 200 μm. Scale bar for C, D, G, H, K, L, O, P is 75 μm. Scale bar for Q, R is 350 μm. Scale bar for S, T is 85 μm. Scale bar for U-X is 100 μm. Abbr. MC - Meckel’s Cartilage, Md – Mandible, Mx- Maxilla, F-Frontal bone, *-Palatine bone in U-X

In contrast, *Med23*^*+/ECKO*^ embryos treated with the HIF1α inhibitor FM19G11, exhibited no change in osteogenesis as measured by Alizarin red staining and mandibular length. However, *Med23*^*ECKO/ECKO*^ mutants subjected to HIF1α inhibition exhibited a marked increase in osteogenesis, particularly in the mandibular and maxillary regions (Fig. 4I-L). This implicates HIF1α-mediated signaling as a key contributor to the impaired ossification observed in *Med23*^*ECKO/ECKO*^ mutants.

The decreased VEGF signaling and increased HIF1α signaling observed in *Med23*^*ECKO/ECKO*^ mutant embryos led us to hypothesize that combinatorial inhibition of HIF1α signaling together with restoration of VEGF signaling could rescue the osteogenic anomalies. We therefore treated pregnant dams with the i-HIF1α+VEGFA at E9.5 and E13.5 and observed a substantial rescue of the craniofacial ossification phenotype in *Med23*^*ECKO/ECKO*^ mutants, as evidenced by restoration of bone formation in the mandible, maxilla, and frontal bones (Fig. 4M-P, U-V, Y).

Furthermore, whereas *Med23*^*ECKO/ECKO*^ mutants are typically embryonic lethal by E16.5, i-HIF1α+VEGFA supplementation prolonged their lifespan for a limited period postnatally. Postnatal day (P) 2 *Med23*^*ECKO/ECKO*^ mutants exhibited no gross structural craniofacial bone defects and were indistinguishable from *Med23*^*+/ECKO*^ controls (Fig. 4Q-T, W-X, Z). However, we were unable to collect pups i-HIF1α+VEGFA supplemented *Med23*^*ECKO/ECKO*^ mutants at P7, for reasons which remain to be determined.

To define the mechanisms underlying the rescue of vascular and osteogenic development, we performed immunostaining on DMSO control and i-HIF1α+VEGFA treated *Med23*^*+/ECKO*^ control and *Med23*^*ECKO/ECKO*^ mutant embryos at E14.5. Runx2 and HIF1α staining revealed restored expression in concert with normal bone size in i-HIF1α+VEGFA treated *Med23*^*ECKO/ECKO*^ mutants comparable to treated *Med23*^*+/ECKO*^ controls, consistent with the rescue of osteogenesis. PECAM1 staining in the mandible revealed a comparable number and distribution of EC in *Med23*^*+/ECKO*^ control and *Med23*^*ECKO/ECKO*^ mutant embryos treated with i-HIF1α+VEGFA (Fig. 5).

**Fig. 5:**
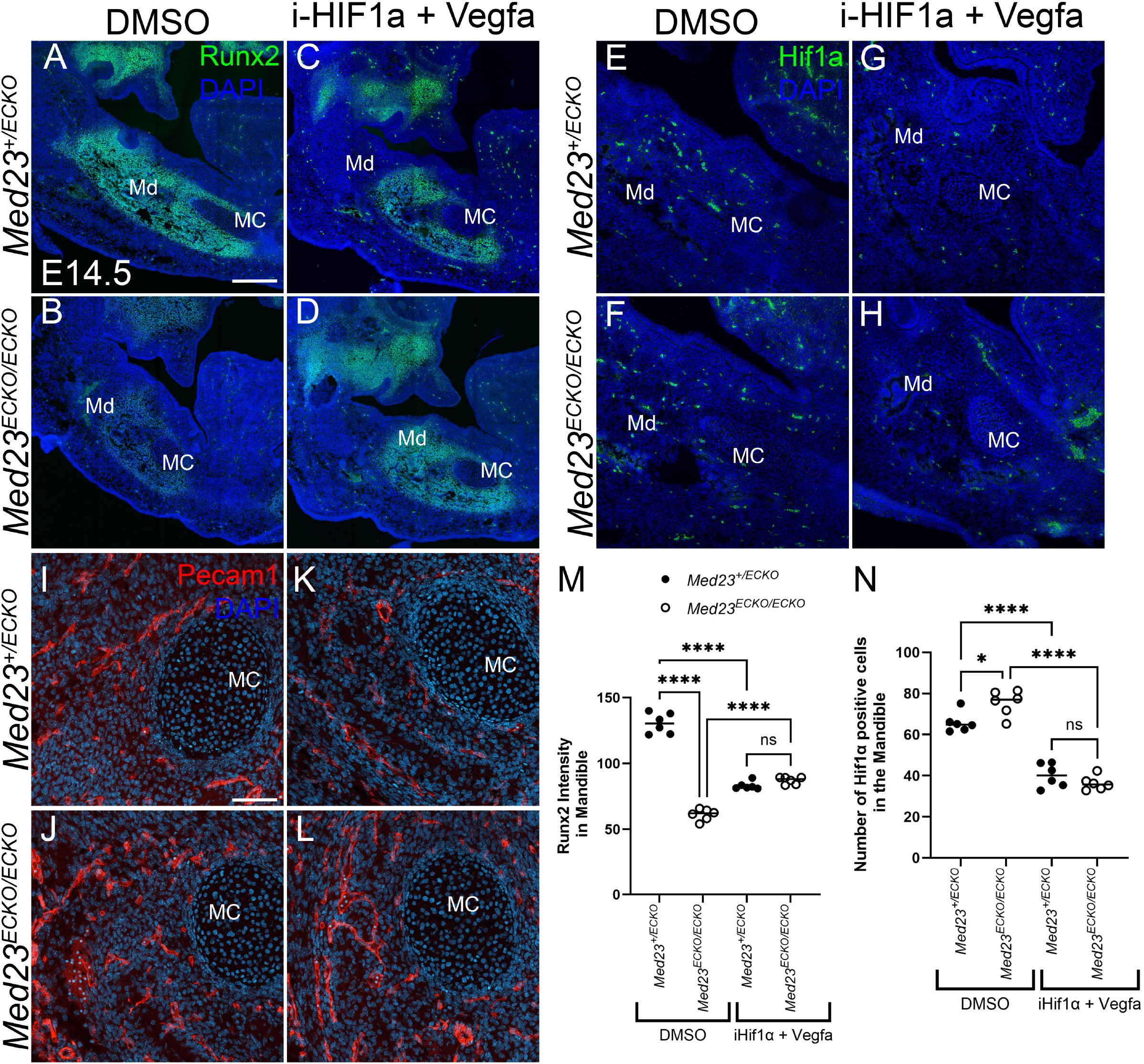
Osteogenesis and angiogenesis rescue of *Med23*^*ECKO/ECKO*^ mutants. *Med23*^*+/ECKO*^ control and *Med23*^*ECKO/ECKO*^ mutant embryos were treated with either DMSO (A, B, E, F, I, J) or a combination of HIF1α inhibitor and recombinant VEGFA (i-HIF1α + Vegfa; C, D, G, H, K, L). Immunostaining was performed for Runx2 (green; A–D), HIF1α (green; E–H), and Pecam1 (red; I-L). Scale bars: 280 μm for panels A–H; 70 μm for panels I-L. (M) Quantification of Runx2 intensity in the mandible (intensity/area). (N) Quantification of HIF1α-positive cells in the mandible. Statistical analysis was performed using one-way ANOVA followed by multiple t-tests.

Together, these findings demonstrate that vascular patterning critically influences craniofacial bone development. Upregulation of HIF1α in *Med23*^*ECKO/ECKO*^ mutants leads to reduced VEGF signaling and impaired ossification, which can be reversed through targeted modulation of HIF1α and VEGF signaling.

## Discussion

In this study, we showed that EC-specific deletion of Med23 results in mid-gestation lethality around E16.5 in association with edema and hemorrhage. Additionally, our study has uncovered a non-cell autonomous effect of ECs and vasculature on embryonic craniofacial bone development. While previous work has implicated Med23 in transcriptional regulation and morphogenesis, our findings extend its function to the regulation of intramembranous ossification via vascular mediated signaling.

NCCs are known to migrate along vascular networks during embryogenesis using endothelial structures as guidance cues (Milgrom-Hoffman et al. 2014). Despite an abnormal vascular network, NCC migration, and differentiation into neurons and chondrocytes in *Med23*^*ECKO/ECKO*^ embryos were normal. However, NCC-derived osteogenesis in the craniofacial region was severely impaired. Although the presence of a cleft palate raises questions regarding potential NCC contributions, our analyses revealed no significant differences in NCC proliferation, apoptosis, or pharyngeal arch morphology, suggesting that the cleft palate phenotype may arise through mechanisms independent of NCC migration or proliferation. We will address these molecular changes in a forthcoming study.

This dissociation between NCC migration and bone formation suggests that EC-derived signals critically guide the differentiation of post-migratory NCCs into osteoblast lineages during craniofacial development. ECs regulate osteogenesis through paracrine signaling, including VEGF and Notch pathways, that support osteoblast maturation and matrix deposition (Ramasamy et al. 2016). In *Med23*^*ECKO/ECKO*^ mutants, disrupted vascular patterning and altered HIF1α signaling likely impair EC-osteoblast communication, reducing expression of key osteogenic markers, such as Runx2 and β-Catenin. These findings support a model in which Med23 functions cell autonomously in EC to maintain vascular integrity and also non-cell autonomously to facilitate the differentiation of NCC-derived osteoprogenitors into craniofacial bone.

The VEGF signaling pathway plays a central role in mediating bone–vascular interactions during development and regeneration. Knockout of Vegfr1 leads to excessive angiogenesis and embryonic lethality, underscoring its critical role in maintaining vascular balance during development (Fong et al. 1999). In bone development, particularly during endochondral ossification, VEGF signaling is regulated by HIF1α to facilitate vascular invasion into a cartilage template, which is essential for osteoblast recruitment and bone matrix deposition (Maes and Carmeliet 2013). However, the relationship between HIF1α-VEGF signaling and osteogenesis is nuanced. While HIF1α promotes Vegfa transcription (Xiao et al. 2021) and osteogenesis, excessive HIF1α activity can inhibit osteoblast differentiation by upregulating Twist2, which suppresses Runx2 (Lee et al. 2024). Interestingly, HIF1α exhibits age-dependent effects: in young bone, it promotes osteogenesis and angiogenesis, whereas in aged bone, elevated HIF1α correlates with impaired bone-vascular coupling, potentially through a ROS-mediated p53 pathway (Shao et al. 2022). This dual role suggests that HIF1α must be tightly regulated to balance angiogenesis and osteogenesis.

Interestingly, our data showed that HIF1α expression was elevated and VEGF signaling was reduced in *Med23*^*ECKO/ECKO*^ mutants, particularly near the mandibular bone. This pattern mirrors the signaling dynamics observed during endochondral ossification, where hypoxia-induced HIF1α promotes osteoblast differentiation through VEGF suppression (Collin-Osdoby 1994). However, in the context of intramembranous ossification, these same molecular cues appear to inhibit bone formation, suggesting a fundamental difference in the vascular dependency of these two ossification processes.

Pharmacological manipulation of HIF1α and VEGF fully restored ossification and extended embryonic viability in *Med23*^*ECKO/ECKO*^ mutants. These findings underscore the importance of tightly regulated hypoxic and angiogenic signaling in craniofacial development and highlight Med23 as a key molecular integrator of these pathways.

In summary, our data show that Med23 is essential for vascular patterning and craniofacial ossification through modulation of HIF1α–VEGF signaling. The differing responses of intramembranous and endochondral ossification to hypoxia and VEGF highlight tissue⍰specific vascular requirements in skeletal development and disease, with implications for improving the repair of critical⍰sized intramembranous craniofacial bone defects.

## Methods

Mice were maintained under standard housing conditions following approved IACUC protocol 23-013 at the University at Albany, SUNY Albany and 2025-184 at the Stowers Institute for Medical Research 2025-184. We have complied with Animal Research: Reporting In Vivo Experiments (ARRIVE) 2.0 checklist.

All other methods and materials are detailed in the appendix.

## Acknowledgements

The authors thank LAS core staff for animal care and husbandry at both SUNY, Albany and the Stowers Institute for Medical Research, especially Melissa Childers and Marina Thexton. This work was funded by K99/R00 (DE030972) from the National Institute for Dental and Craniofacial Research (S.D.) and the Stowers Institute for Medical Research (P.A.T). Part of the research reported in this publication was supported by the Office of the Director, National Institutes of Health under Award Number S10OD028600 and S10OD036207. The authors do not have any conflicts of interest. Original data is located at http://www.stowers.org/research/publications/LIBPB-2590 and https://doi.org/10.7910/DVN/I0QP2C. R code for Spatial analysis can be found at https://github.com/mmarchin/cbio.sdash.101.

## Author contributions

S Dash: Contributed to conception, design, data acquisition and interpretation, drafted and critically revised the manuscript

J R Rettig: Contributed to data acquisition and interpretation, drafted and critically revised the manuscript

M. Gogol: Contributed to data acquisition and interpretation, critically revised the manuscript P A Trainor: Contributed to conception, design, and critically revised the manuscript

All authors gave their final approval and agreed to be accountable for all aspects of the work.

