## Supplementary material for "Vascular Patterning Shapes Intramembranous Ossification via HIF1α-VEGF Axis": Supplemetnary 1

**Appendix file**

**Appendix materials and methods:**

**Mouse husbandry**

### *Med23flox* animals were obtained from KOMP and maintained as previously described (Dash et al. 2020). Similarly, *Tek-Cre* (Tie2-Cre) (B6.Cg-Tg(Tek-Cre)1Ywa/J) and *Rosa-eYFP* (B6.129X1-*Gt(ROSA)26Sor^tm1(EYFP)Cos^*/J) mice were obtained from JAX and maintained as previously described (Dash et al. 2020; Dash et al. 2021). For timed matings, the day a vaginal plug was found was designated as embryonic day 0.5 (E0.5). *Med23^fx/fx^;Tek-Cre* (*Med23^ECKO/ECKO^*) and *Med23^+/fx^;Tek-Cre* (*Med23^+/ECKO^*) embryos were used as mutants and controls for every experiment, respectively.

**Brightfield imaging and Skeletal staining**

Embryos were dissected at the desired stages in PBS, anesthetized in ice cold PBS for an hour and imaged using a Nikon DS-Ri camera or a Nikon DS10 Digital Camera. For skeletal preparation and staining, embryos were collected and fixed in 95% ethanol, followed by Alcian blue and Alizarin red staining of cartilage and bone as previously described (Behringer et al. 2014). Stained embryos were imaged using a Nikon DS-Ri camera or a Nikon DS10 Digital Camera. For quantification, bone length was measured in ImageJ.

**Histology**

Embryos were dissected at the desired developmental stages in PBS and fixed in 4% paraformaldehyde, then dehydrated to 70% ethanol for paraffin embedding. Paraffin‑embedded embryos were sectioned at 5 µm thickness and stained with hematoxylin and eosin using standard protocols. Stained slides were imaged using an EVIDENT VS200 slide scanner.

**Lineage tracing**

*Med23^fx/+^;Tek-Cre* mice were crossed with *Rosa-eYFP* mice to generate *Med23^fx/+^;Tek-Cre;YFP+* and *Med23^fx/fx^;Tek-Cre;YFP+* embryos. These animals were then immunostained with a GFP antibody as described below. Quantification of cranial blood‑vessel dimensions was performed as follows: for vessel width, we measured the widest point of each vessel within an embryo and calculated the average width across all vessels. For vessel length, we measured the distance from the vessel root to the distal tip as visualized by GFP staining and averaged these lengths for each embryo.

**Immunostaining**

Whole embryo and dissected palates were fixed in 4% paraformaldehyde overnight followed by dehydration through an ascending methanol series into 100% methanol, after which they were stored at -20C until further use as described previously (Dash et al. 2021). The tissues were then treated with Dent’s bleach (4:1:1 Methanol : DMSO : Hydrogen peroxide), rehydrated, and processed for immunostaining as previously described (Dash et al. 2021; Falcon et al. 2021).

Tissue sections both FFPE and frozen were stained according to standard protocols. Antibodies used were: Pecam1 (CD31, 1:100, BD Pharmingen #553370), Cleaved Caspase 3 (1:100, Cell signaling #9661S), Phospho-Histone H3 (1:1000, Millipore #06-570), Sox9 (1:500, Abcam #ab185966), Sox10 (1:500, Abcam #ab155279), Tuj1(1:1000, Biolegend #657402), Runx2 (1:500, Abcam #ab192256), b-Catenin (1:500, Abcam #ab32572) and HIF1α (1:1000, Abcam #ab228649). Each experiment consisted of 5 biological replicates and 3 technical replicates. ImageJ was used for analysis and quantification. For quantification, technical replicates were averaged per sample. Intensity measurements were normalized to the area.

Sox9 and Sox10 quantifications were performed specifically on pharyngeal arch 1 and trigeminal ganglion, respectively. Volumetric measurements were performed using IMARIS.

**Spatial Transcriptomics**

E14.5 *Med23^+/ECKO^* and *Med23^ECKO/ECKO^* embryos were dissected in PBS, fixed in 4% paraformaldehyde and then embedded in paraffin. The samples were sectioned serially at 10um. One of the slides with serial sections was stained with Hematoxylin and Eosin, whereas other slides were used for spatial transcriptomics with the NanoString COSMX platform. The experiment was performed using two technical replicates and two biological replicates. Analysis of Spatial Transcriptomics was performed using Seurat Package (4.9.9.9041) in R (4.2.0).

**qPCR**

Dissected mandibles from embryos treated with HIF1α inhibitor, FM19G11 (5mg/kg, Selleckchem #E1712), HIF1α activator, DMOG (3mg/kg, Selleckchem #S7483) and DMSO were used for RNA isolation. A total of 5 *Med23^+/ECKO^* and 5 *Med23^ECKO/ECKO^* embryos were used for the experiment. RNA isolation was performed using RNeasy Plus Mini kit. The Superscript III Kit (Invitrogen) was used to synthesize cDNA for quantitative RT-PCR (qPCR) using random hexamer primers. qPCR was performed on ABI7000 (Thermo QuantStudio 7) using Perfecta Sybr Green (Quantbio # 95072-250).

Primers are as follows:

| Gene | Forward | Reverse |
| --- | --- | --- |
| Col1a1 | GTATGCTTGATCTGTATCTGC | TGGGTCCCGTATTCTTCCG |
| Col10a1 | CATCTCCCAGCACCAGAATC | CCTGGCAAGCCTTGTTTTCC |
| Runx2 | GCGGACGAGGCAAGAGTT | CGTGTGGAAGACAGCGGC |
| Hif1a | TTCAAGCAGCAGGAATTGGAAC | TGAATGTGCTGTGATCTGGCA |
| Spp1 | TTGCTTTTGCCTGTTTGGCA | GTGCAGGCTGTAAAGCTTCTT |

**Drug Treatments for Phenotype Rescue**

Pregnant dams were treated with the drugs mentioned below via intraperitoneal injection once at E9.5 and again at E13.5. Drugs included HIF1α inhibitor, FM19G11 (20mg/kg, Selleckchem #E1712), HIF1α activator, DMOG (10mg/kg, Selleckchem #S7483), VEGF Signaling inhibitor, ZM323881 (10mg/kg, Selleckchem # S2896) and VEGF signaling activator, VEGFA (50ug/kg, Abnova, # P4608). DMSO was used as a vehicle control. The embryos were collected at E14.5 and E16.5 and neonatal pups were collected at P2 and P7 for end point experiments. Drugs used included HIF1α inhibitor, FM19G11 (5mg/kg, Selleckchem #E1712), HIF1α activator, DMOG (3mg/kg, Selleckchem #S7483). A total of 5 *Med23^+/ECKO^* and 5 *Med23^ECKO/ECKO^* embryos were used for the experiment.

**Statistical tests**

Student t-tests and ANOVA with multiple t-tests were performed based on the number of numerical variables and comparisons. When applicable non-parametric Mann-Whitney U-tests were performed for multiple comparisons. P value of <0.05 was designated with one asterisk *, <0.005 with ** , <0.0001 with *** and <0.00001 with ****.

**Appendix Figures**

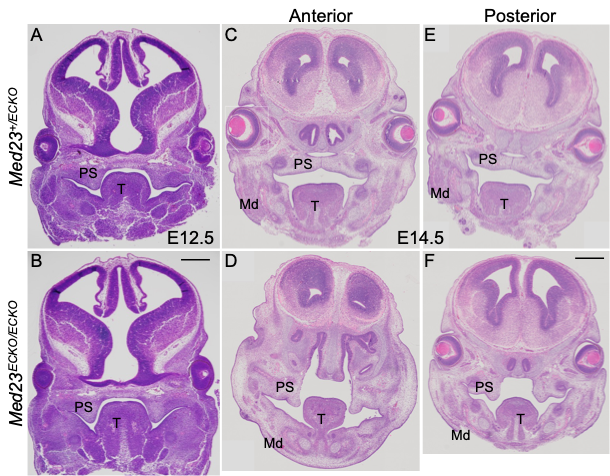
**Appendix Figure 1: Histological analysis of *Med23^+/ECKO^* control and *Med23^ECKO/ECKO^* mutant embryos.** (A-B) Hematoxylin and Eosin stained sections at E12.5 show no detectable differences between control and mutant embryos in the size or tissue organization of the palatal shelves or the lower jaw. (C–F) By E14.5, the anterior and posterior palatal shelves are fully fused in control embryos, whereas in *Med23^ECKO/ECKO^* mutants the shelves remain vertically oriented. *Med23^ECKO/ECKO^* mutant embryos also exhibit a reduction in mandibular size at this stage.

Scale bar for A and B is 100 μm and C-F is 170 μm.

Abbr. PS: Palatal Shelf, T, Tongue, Md- Mandible

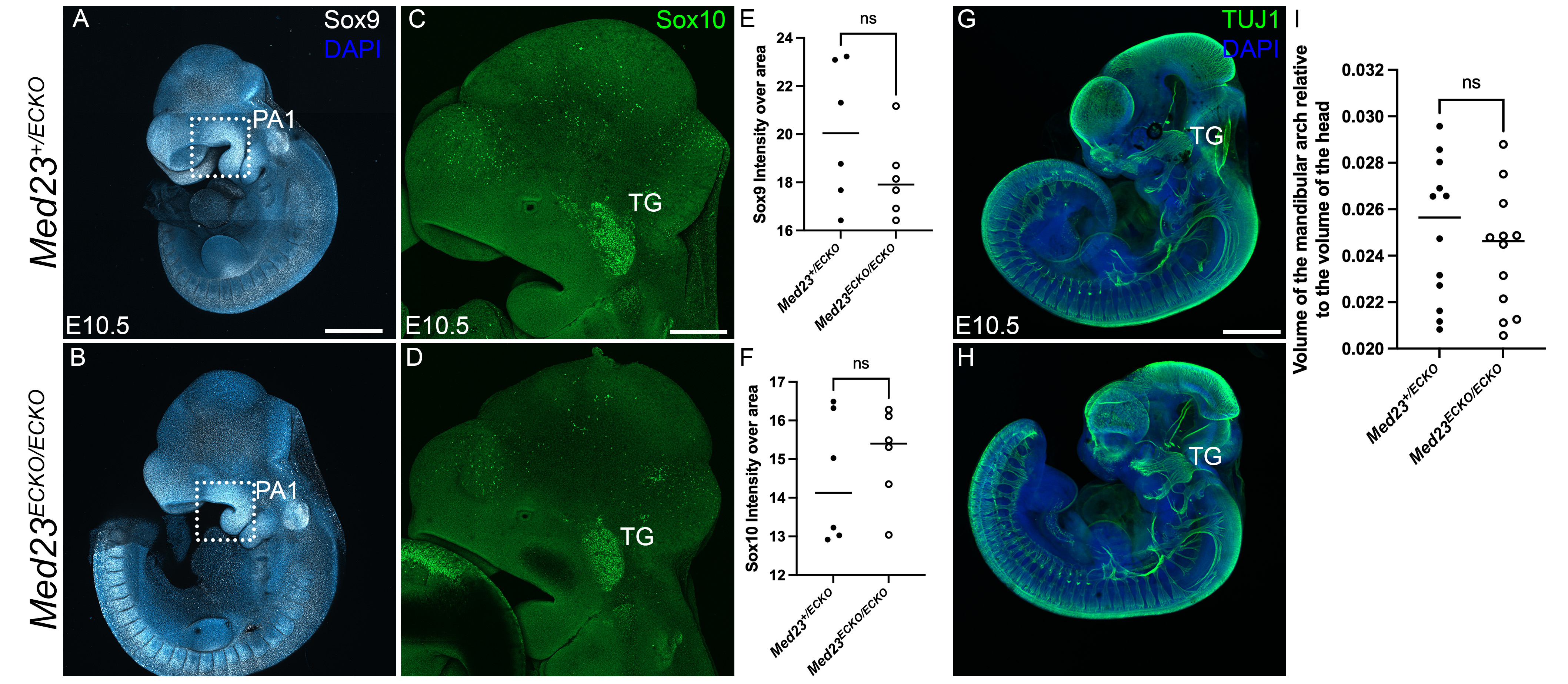

**Appendix Figure 2: Neural crest cell migration and differentiation into neuronal lineages is not affected in *Med23^ECKO/ECKO^* mutant embryos.** Immunostaining for neural crest cell markers, Sox9 (A,B,E, scale bar = 250 μm) and Sox10 at E10.5 (C,D,F, scale bar = 150 μm) suggests neural crest cells and neuronal cell progenitors are unaffected in the *Med23^ECKO/ECKO^* embryos (n=6)*.* (G,H) Neuronal cells labeled by TUJ1 are similar between *Med23^+/ECKO^* control and *Med23^ECKO/ECKO^* mutant embryos (n=3). Scale bar = 250 μm. (I) Volumetric measurement of the mandibular arch relative to the head indicates no significant difference between *Med23^+/ECKO^* control and *Med23^ECKO/ECKO^* mutant embryos (n=12). Abbr. PA1: Pharyngeal arch 1, TG: Trigeminal ganglion

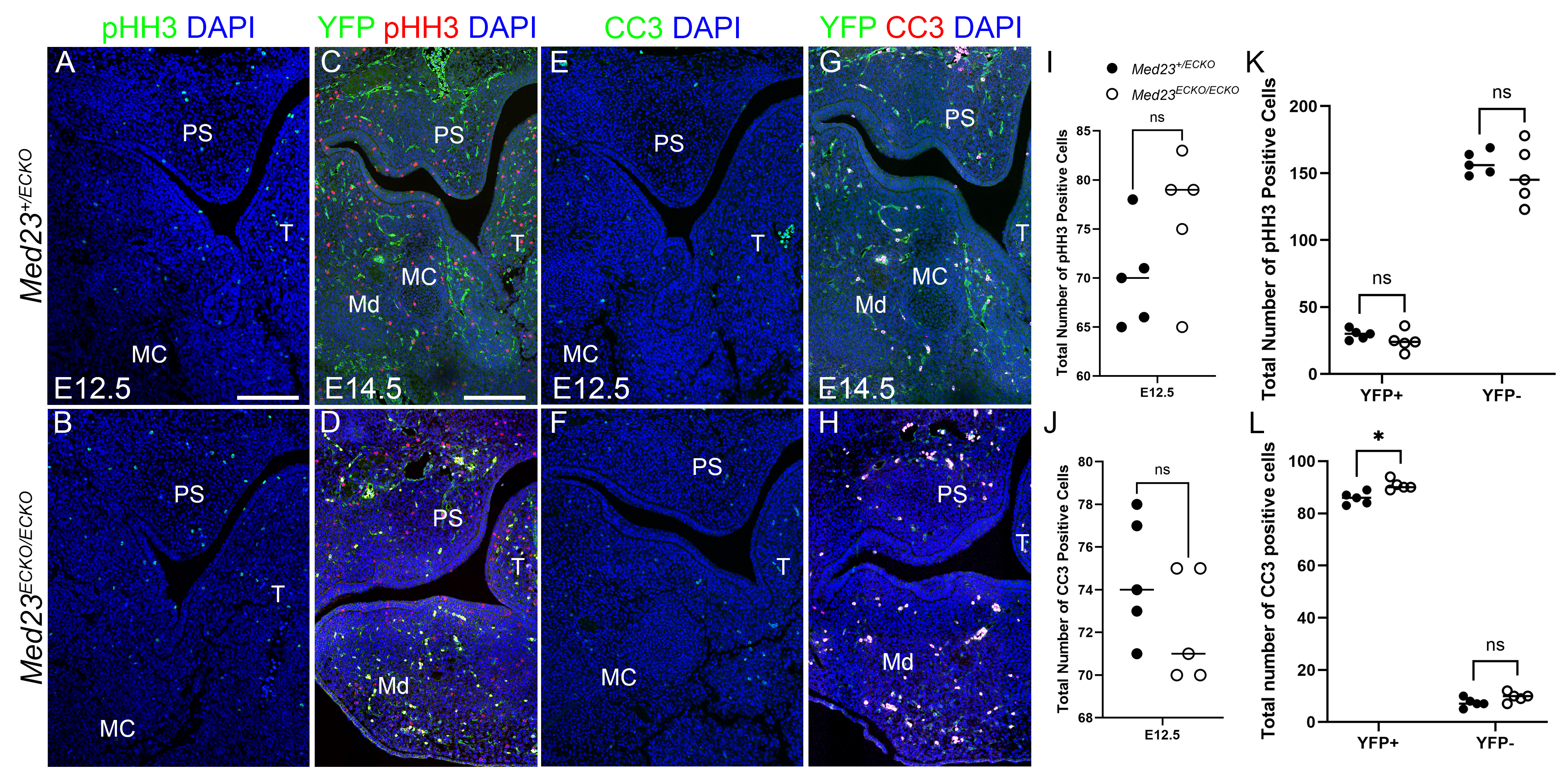
**Appendix Figure 3: Proliferation and apoptosis anomalies of *Med23^ECKO/ECKO^* mutants.** Section immunostaining of *Med23^+/ECKO^* controls and *Med23^ECKO/ECKO^* mutants was performed with phospho-histone H3 (pHH3, green in A,B, red in C,D) and Cleaved Caspase 3 (CC3, green in E,F, red in G,H) at E12.5 and E14.5. E14.5 sections were also stained with YFP to lineage trace endothelial cells (C, D, G, H). n = 5. Scale bar is 70 μm. Quantifications of pHH3 and CC3 positive cells at E12.5 are shown in I and J, respectively. Quantifications of pHH3 and CC3 positive cells at E14.5 in the YFP+ (Endothelial cell) population vs YFP- cells are shown in K and L, respectively. Abbr. PS - Palatal Shelf, T – Tongue, MC - Meckel’s Cartilage, Md – Mandible

**Appendix Table 1:** List of differentially expressed genes identified through spatial transcriptomics. Genes are listed with cluster numbers.

| **gene** | **p_val** | **avg_log2FC** | **pct.1** | **pct.2** | **p_val_adj** | **Cluster number** |
| --- | --- | --- | --- | --- | --- | --- |
| Gnas | 0 | -0.5178197 | 0.789 | 0.877 | 0 | 0 |
| Aldoa | 1.41E-175 | 0.55745425 | 0.496 | 0.356 | 1.36E-172 | 0 |
| Tuba1a/b/c | 2.16E-127 | 0.26735281 | 0.907 | 0.841 | 2.07E-124 | 0 |
| Col3a1 | 4.10E-113 | -0.3196454 | 0.762 | 0.812 | 3.93E-110 | 0 |
| Ndrg1 | 3.83E-81 | 0.5036035 | 0.152 | 0.087 | 3.67E-78 | 0 |
| Grb10 | 1.97E-79 | -0.3623163 | 0.594 | 0.648 | 1.89E-76 | 0 |
| Hspa8 | 2.25E-79 | -0.3617872 | 0.517 | 0.588 | 2.16E-76 | 0 |
| Cd9 | 1.10E-68 | 0.43421751 | 0.149 | 0.088 | 1.05E-65 | 0 |
| Gnai2 | 2.63E-67 | 0.32378828 | 0.394 | 0.305 | 2.52E-64 | 0 |
| Ptn | 6.66E-67 | -0.3531218 | 0.519 | 0.577 | 6.39E-64 | 0 |
| Smad2 | 5.32E-64 | 0.3962261 | 0.176 | 0.112 | 5.11E-61 | 0 |
| P2rx7 | 6.61E-61 | 0.42816219 | 0.115 | 0.065 | 6.34E-58 | 0 |
| Bin1 | 1.43E-57 | 0.36116968 | 0.233 | 0.165 | 1.37E-54 | 0 |
| Tcf12 | 5.09E-57 | 0.32900454 | 0.286 | 0.211 | 4.89E-54 | 0 |
| Maged1 | 3.44E-55 | -0.2558238 | 0.65 | 0.693 | 3.30E-52 | 0 |
| Atpif1 | 2.23E-54 | 0.29926307 | 0.357 | 0.279 | 2.13E-51 | 0 |
| Adgrl2 | 4.41E-54 | 0.38294397 | 0.19 | 0.13 | 4.23E-51 | 0 |
| Dnm1 | 5.74E-51 | 0.30775015 | 0.318 | 0.245 | 5.51E-48 | 0 |
| Anapc16 | 1.61E-50 | 0.36850449 | 0.19 | 0.131 | 1.54E-47 | 0 |
| Serpine2 | 3.52E-50 | 0.32837248 | 0.234 | 0.17 | 3.37E-47 | 0 |
| Pcsk6 | 3.37E-49 | 0.38110466 | 0.145 | 0.094 | 3.23E-46 | 0 |
| Gpr37l1 | 4.35E-49 | 0.36780101 | 0.103 | 0.06 | 4.17E-46 | 0 |
| Mapk14 | 4.94E-49 | 0.34602828 | 0.187 | 0.129 | 4.74E-46 | 0 |
| Atr | 9.19E-49 | 0.37565851 | 0.124 | 0.077 | 8.82E-46 | 0 |
| Bcl2 | 6.79E-48 | 0.36917143 | 0.159 | 0.106 | 6.52E-45 | 0 |
| Mib1 | 8.92E-48 | 0.33670985 | 0.195 | 0.137 | 8.55E-45 | 0 |
| Sstr4 | 2.02E-47 | 0.36046946 | 0.123 | 0.077 | 1.94E-44 | 0 |
| Mid1 | 8.38E-47 | 0.34611034 | 0.216 | 0.157 | 8.04E-44 | 0 |
| Bsg | 1.62E-46 | 0.2871058 | 0.309 | 0.239 | 1.55E-43 | 0 |
| Mef2a | 1.92E-46 | 0.35215834 | 0.143 | 0.094 | 1.84E-43 | 0 |
| Vegfa | 2.49E-46 | 0.31775813 | 0.252 | 0.189 | 2.38E-43 | 0 |
| Lgr5 | 6.90E-45 | 0.35875854 | 0.117 | 0.073 | 6.62E-42 | 0 |
| Mmp2 | 1.43E-44 | -0.3988477 | 0.276 | 0.335 | 1.37E-41 | 0 |
| Sv2a | 5.10E-44 | 0.35830064 | 0.128 | 0.083 | 4.89E-41 | 0 |
| Itm2a | 7.28E-44 | -0.3763202 | 0.386 | 0.441 | 6.98E-41 | 0 |
| Ngfr | 1.20E-43 | 0.3011833 | 0.15 | 0.1 | 1.15E-40 | 0 |
| Hipk2 | 6.52E-43 | 0.34185839 | 0.154 | 0.105 | 6.25E-40 | 0 |
| Pten | 9.34E-43 | 0.30504716 | 0.196 | 0.141 | 8.96E-40 | 0 |
| Tnrc6a | 1.02E-42 | 0.33123604 | 0.15 | 0.102 | 9.82E-40 | 0 |
| Smurf1 | 3.69E-42 | 0.33979906 | 0.143 | 0.096 | 3.54E-39 | 0 |
| Axl | 7.55E-41 | 0.29278314 | 0.217 | 0.16 | 7.24E-38 | 0 |
| Cox4i1 | 2.05E-40 | 0.25458594 | 0.379 | 0.311 | 1.96E-37 | 0 |
| Lifr | 2.50E-40 | 0.32489937 | 0.111 | 0.07 | 2.39E-37 | 0 |
| Calm1 | 1.74E-39 | -0.2740641 | 0.513 | 0.557 | 1.67E-36 | 0 |
| Ghr | 2.21E-39 | 0.33238759 | 0.139 | 0.094 | 2.12E-36 | 0 |
| Stk3 | 4.74E-39 | 0.29568806 | 0.141 | 0.095 | 4.55E-36 | 0 |
| Tgfb1 | 5.12E-39 | 0.32782037 | 0.108 | 0.069 | 4.91E-36 | 0 |
| Cd55 | 1.76E-38 | 0.33369734 | 0.123 | 0.081 | 1.69E-35 | 0 |
| Ctnnb1 | 4.05E-38 | -0.3280599 | 0.236 | 0.293 | 3.88E-35 | 0 |
| Fth1 | 8.34E-38 | -0.253062 | 0.56 | 0.599 | 8.00E-35 | 0 |
| Mertk | 1.00E-37 | 0.29289762 | 0.16 | 0.113 | 9.60E-35 | 0 |
| Wac | 1.15E-37 | 0.27057407 | 0.262 | 0.203 | 1.10E-34 | 0 |
| Thy1 | 1.20E-37 | 0.33982321 | 0.151 | 0.105 | 1.15E-34 | 0 |
| Ctxn1 | 1.92E-37 | 0.29158217 | 0.217 | 0.163 | 1.84E-34 | 0 |
| Epb41l2 | 9.65E-37 | 0.28806609 | 0.229 | 0.175 | 9.26E-34 | 0 |
| Nsg1 | 3.10E-36 | 0.2639404 | 0.252 | 0.195 | 2.97E-33 | 0 |
| Dnm1l | 3.28E-36 | 0.29173804 | 0.183 | 0.133 | 3.15E-33 | 0 |
| Ndufa4 | 3.46E-36 | -0.3235726 | 0.281 | 0.336 | 3.32E-33 | 0 |
| Sparc | 4.20E-36 | -0.2540146 | 0.523 | 0.569 | 4.03E-33 | 0 |
| Ldha | 1.72E-35 | 0.25250949 | 0.429 | 0.366 | 1.65E-32 | 0 |
| Samhd1 | 2.89E-35 | 0.3054553 | 0.183 | 0.134 | 2.77E-32 | 0 |
| Trp53 | 7.21E-35 | 0.29596105 | 0.176 | 0.128 | 6.91E-32 | 0 |
| Pde8a | 1.15E-34 | 0.296538 | 0.1 | 0.064 | 1.11E-31 | 0 |
| Wdr59 | 2.27E-34 | 0.29899883 | 0.141 | 0.099 | 2.17E-31 | 0 |
| Bmpr2 | 2.75E-34 | 0.27457061 | 0.213 | 0.162 | 2.64E-31 | 0 |
| Creb1 | 5.17E-34 | 0.28712155 | 0.192 | 0.143 | 4.96E-31 | 0 |
| Cck | 1.76E-33 | 0.29517193 | 0.117 | 0.078 | 1.68E-30 | 0 |
| Rab1a | 4.67E-33 | -0.3209591 | 0.301 | 0.351 | 4.48E-30 | 0 |
| Atp6v0a1 | 5.76E-33 | 0.29977231 | 0.11 | 0.073 | 5.52E-30 | 0 |
| Slc11a2 | 1.78E-32 | 0.29570085 | 0.159 | 0.115 | 1.70E-29 | 0 |
| Phc2 | 3.40E-32 | 0.25271705 | 0.229 | 0.177 | 3.26E-29 | 0 |
| Apoe | 2.21E-31 | 0.25517155 | 0.152 | 0.11 | 2.12E-28 | 0 |
| Cfh | 2.22E-31 | 0.2769407 | 0.118 | 0.081 | 2.13E-28 | 0 |
| Cspg5 | 2.74E-31 | 0.27960206 | 0.106 | 0.071 | 2.63E-28 | 0 |
| Btrc | 7.93E-31 | 0.30423201 | 0.123 | 0.086 | 7.61E-28 | 0 |
| Tagln | 1.15E-29 | 0.28480737 | 0.113 | 0.077 | 1.10E-26 | 0 |
| Usp15 | 4.39E-29 | 0.2766911 | 0.157 | 0.116 | 4.21E-26 | 0 |
| Lpar2 | 4.59E-29 | 0.28035779 | 0.117 | 0.081 | 4.40E-26 | 0 |
| Lrp6 | 1.27E-28 | 0.25430908 | 0.163 | 0.121 | 1.22E-25 | 0 |
| Lilrb4a/b | 2.23E-28 | 0.28774474 | 0.136 | 0.098 | 2.14E-25 | 0 |
| Ndufs1 | 5.50E-28 | 0.25328181 | 0.199 | 0.154 | 5.27E-25 | 0 |
| Psen1 | 8.10E-28 | 0.25693407 | 0.116 | 0.081 | 7.77E-25 | 0 |
| Jak2 | 1.68E-27 | 0.27282513 | 0.16 | 0.119 | 1.61E-24 | 0 |
| Clu | 4.38E-26 | 0.26148157 | 0.132 | 0.096 | 4.20E-23 | 0 |
| Pak1 | 6.31E-26 | 0.25948388 | 0.145 | 0.108 | 6.05E-23 | 0 |
| Hspa8 | 1.06E-100 | -0.59727 | 0.616 | 0.77 | 1.02E-97 | 1 |
| Tmsb10 | 2.20E-100 | -0.3921636 | 0.979 | 0.991 | 2.11E-97 | 1 |
| Gnas | 3.04E-89 | -0.3959091 | 0.895 | 0.951 | 2.91E-86 | 1 |
| Acta2 | 4.84E-72 | -0.5056794 | 0.749 | 0.838 | 4.64E-69 | 1 |
| Aldoa | 5.90E-42 | 0.36410899 | 0.791 | 0.693 | 5.66E-39 | 1 |
| Grb10 | 3.66E-30 | -0.295842 | 0.719 | 0.786 | 3.51E-27 | 1 |
| Fth1 | 6.90E-29 | -0.3455912 | 0.513 | 0.618 | 6.62E-26 | 1 |
| Ctnnb1 | 6.92E-29 | -0.5061111 | 0.202 | 0.308 | 6.64E-26 | 1 |
| Ndufa4 | 1.09E-23 | -0.357419 | 0.399 | 0.498 | 1.05E-20 | 1 |
| Vegfa | 1.30E-18 | 0.40582409 | 0.236 | 0.164 | 1.24E-15 | 1 |
| Ldha | 3.19E-18 | 0.32088082 | 0.61 | 0.546 | 3.06E-15 | 1 |
| Atp2a2 | 7.19E-17 | -0.3256572 | 0.405 | 0.484 | 6.89E-14 | 1 |
| Calm2 | 1.07E-16 | -0.2509092 | 0.611 | 0.669 | 1.03E-13 | 1 |
| Rab1a | 2.11E-16 | -0.3704751 | 0.264 | 0.343 | 2.02E-13 | 1 |
| Gnai2 | 2.49E-16 | 0.36012842 | 0.355 | 0.279 | 2.39E-13 | 1 |
| Mid1 | 2.60E-16 | 0.44365991 | 0.267 | 0.2 | 2.49E-13 | 1 |
| Sod1 | 4.09E-14 | -0.2587705 | 0.536 | 0.601 | 3.93E-11 | 1 |
| Gdnf | 5.91E-14 | 0.37458484 | 0.102 | 0.06 | 5.66E-11 | 1 |
| Mertk | 1.66E-13 | 0.3562195 | 0.124 | 0.078 | 1.60E-10 | 1 |
| Atr | 3.37E-13 | 0.32116621 | 0.151 | 0.101 | 3.24E-10 | 1 |
| Idnk | 8.35E-13 | 0.36497489 | 0.106 | 0.066 | 8.00E-10 | 1 |
| Tcf12 | 2.16E-12 | 0.2737625 | 0.349 | 0.28 | 2.07E-09 | 1 |
| Anapc16 | 1.22E-11 | 0.30115052 | 0.206 | 0.152 | 1.17E-08 | 1 |
| Atp2b1 | 2.23E-11 | -0.2829436 | 0.27 | 0.335 | 2.14E-08 | 1 |
| Epb41l3 | 4.07E-11 | 0.29088703 | 0.258 | 0.2 | 3.90E-08 | 1 |
| Flt1 | 4.10E-11 | 0.34692205 | 0.1 | 0.063 | 3.93E-08 | 1 |
| Psmd1 | 4.99E-11 | -0.3418781 | 0.167 | 0.222 | 4.78E-08 | 1 |
| Pfkl | 7.12E-11 | 0.30521992 | 0.269 | 0.212 | 6.83E-08 | 1 |
| Cd55 | 7.69E-11 | 0.30653145 | 0.129 | 0.088 | 7.38E-08 | 1 |
| Ttc3 | 8.26E-11 | -0.2769602 | 0.293 | 0.356 | 7.92E-08 | 1 |
| Cd9 | 8.38E-11 | 0.30171928 | 0.128 | 0.086 | 8.04E-08 | 1 |
| Slc17a7 | 1.36E-10 | 0.29246973 | 0.112 | 0.074 | 1.31E-07 | 1 |
| P2rx7 | 2.36E-10 | 0.31791794 | 0.111 | 0.073 | 2.26E-07 | 1 |
| Usp15 | 2.69E-10 | 0.28252915 | 0.234 | 0.18 | 2.58E-07 | 1 |
| Tjp1 | 2.92E-10 | 0.30246581 | 0.173 | 0.127 | 2.80E-07 | 1 |
| Des | 5.04E-10 | -0.2568842 | 0.467 | 0.518 | 4.83E-07 | 1 |
| Smad2 | 5.61E-10 | 0.31455315 | 0.189 | 0.142 | 5.38E-07 | 1 |
| Agtr2 | 1.17E-09 | -0.2607311 | 0.279 | 0.338 | 1.12E-06 | 1 |
| Rgs6 | 1.58E-09 | 0.27464496 | 0.109 | 0.073 | 1.51E-06 | 1 |
| Dnm1 | 2.28E-09 | 0.29045613 | 0.188 | 0.142 | 2.18E-06 | 1 |
| Smurf1 | 2.45E-09 | 0.30813338 | 0.168 | 0.124 | 2.35E-06 | 1 |
| Atp6v0a1 | 5.33E-09 | 0.25623422 | 0.18 | 0.135 | 5.12E-06 | 1 |
| Lifr | 8.09E-09 | 0.27260439 | 0.153 | 0.112 | 7.76E-06 | 1 |
| Fxyd6 | 8.58E-09 | -0.2970823 | 0.19 | 0.24 | 8.23E-06 | 1 |
| Trim2 | 9.27E-09 | 0.26886504 | 0.128 | 0.091 | 8.89E-06 | 1 |
| Mib1 | 1.23E-08 | 0.26050439 | 0.21 | 0.164 | 1.18E-05 | 1 |
| Tcf7l2 | 1.72E-08 | -0.2576036 | 0.315 | 0.368 | 1.65E-05 | 1 |
| Ctxn1 | 1.74E-08 | 0.31186001 | 0.198 | 0.155 | 1.67E-05 | 1 |
| Btrc | 3.01E-08 | 0.29688766 | 0.154 | 0.115 | 2.88E-05 | 1 |
| Ntrk1 | 5.86E-08 | 0.26346401 | 0.163 | 0.124 | 5.62E-05 | 1 |
| Trp53 | 6.39E-08 | 0.26195611 | 0.186 | 0.144 | 6.13E-05 | 1 |
| Pik3cb | 6.92E-08 | 0.25715437 | 0.102 | 0.07 | 6.64E-05 | 1 |
| Ndrg1 | 8.27E-08 | 0.25839182 | 0.151 | 0.113 | 7.93E-05 | 1 |
| Lpar2 | 8.32E-08 | 0.2727549 | 0.119 | 0.086 | 7.98E-05 | 1 |
| Cbln2 | 1.30E-07 | 0.2502151 | 0.123 | 0.089 | 0.00012486 | 1 |
| Lilrb4a/b | 2.45E-07 | 0.26500476 | 0.151 | 0.114 | 0.00023464 | 1 |
| Pak1 | 3.44E-07 | 0.27155866 | 0.123 | 0.091 | 0.00033003 | 1 |
| Sox6 | 7.76E-07 | 0.25999692 | 0.132 | 0.099 | 0.00074454 | 1 |
| Stk3 | 1.20E-06 | 0.2605763 | 0.148 | 0.114 | 0.00115111 | 1 |
| Cacna1b | 1.59E-06 | 0.25181974 | 0.12 | 0.09 | 0.00152826 | 1 |
| Sv2a | 5.35E-06 | 0.25544833 | 0.119 | 0.091 | 0.00512991 | 1 |
| Slco3a1 | 1.69E-05 | 0.25373069 | 0.124 | 0.096 | 0.01618755 | 1 |
| Cxcl14 | 3.13E-05 | -0.2617818 | 0.09 | 0.117 | 0.03004828 | 1 |
| Col1a1 | 5.11E-85 | 0.42575366 | 0.915 | 0.828 | 4.90E-82 | 2 |
| Gnas | 1.96E-74 | -0.3876552 | 0.892 | 0.935 | 1.88E-71 | 2 |
| Aldoa | 2.57E-69 | 0.62517681 | 0.522 | 0.358 | 2.46E-66 | 2 |
| Adgrl2 | 2.98E-48 | 0.62529447 | 0.307 | 0.187 | 2.86E-45 | 2 |
| Hspa8 | 1.91E-42 | -0.4729589 | 0.474 | 0.578 | 1.83E-39 | 2 |
| Dnm1 | 9.92E-41 | 0.46677498 | 0.395 | 0.269 | 9.51E-38 | 2 |
| Bcl2 | 2.89E-35 | 0.55842623 | 0.191 | 0.106 | 2.78E-32 | 2 |
| Timp2 | 1.19E-31 | 0.42854183 | 0.403 | 0.295 | 1.14E-28 | 2 |
| Epb41l2 | 1.73E-28 | 0.44122124 | 0.268 | 0.177 | 1.66E-25 | 2 |
| Serpine2 | 1.06E-26 | 0.47329616 | 0.212 | 0.133 | 1.01E-23 | 2 |
| Ldha | 6.18E-23 | 0.38991323 | 0.378 | 0.29 | 5.93E-20 | 2 |
| Tcf12 | 1.76E-22 | 0.36942153 | 0.326 | 0.24 | 1.69E-19 | 2 |
| Tcf7l2 | 2.45E-22 | -0.4300331 | 0.387 | 0.465 | 2.35E-19 | 2 |
| Selenow | 7.43E-22 | 0.35039211 | 0.363 | 0.275 | 7.13E-19 | 2 |
| Atp1b1 | 1.86E-21 | 0.41556904 | 0.215 | 0.144 | 1.79E-18 | 2 |
| Atpif1 | 2.52E-20 | 0.33155634 | 0.351 | 0.265 | 2.41E-17 | 2 |
| Prnp | 4.91E-20 | 0.43047759 | 0.127 | 0.074 | 4.71E-17 | 2 |
| Gap43 | 9.30E-20 | 0.35735871 | 0.3 | 0.221 | 8.92E-17 | 2 |
| Bsg | 8.22E-19 | 0.30891615 | 0.339 | 0.256 | 7.88E-16 | 2 |
| Lrp6 | 9.14E-19 | 0.38407236 | 0.205 | 0.139 | 8.76E-16 | 2 |
| Mertk | 1.38E-18 | 0.38401267 | 0.147 | 0.091 | 1.32E-15 | 2 |
| Ndufa4 | 6.60E-18 | -0.373784 | 0.291 | 0.363 | 6.33E-15 | 2 |
| Gnai2 | 1.03E-17 | 0.31357505 | 0.375 | 0.295 | 9.85E-15 | 2 |
| Creb1 | 1.55E-17 | 0.34913291 | 0.209 | 0.144 | 1.49E-14 | 2 |
| Ndrg1 | 3.38E-17 | 0.40966793 | 0.164 | 0.108 | 3.25E-14 | 2 |
| Cox4i1 | 3.77E-17 | 0.32396559 | 0.395 | 0.318 | 3.61E-14 | 2 |
| Cd55 | 6.44E-17 | 0.39237008 | 0.154 | 0.1 | 6.18E-14 | 2 |
| Camk1d | 7.85E-17 | 0.3666609 | 0.1 | 0.057 | 7.53E-14 | 2 |
| Pde8a | 9.98E-17 | 0.39035082 | 0.102 | 0.058 | 9.57E-14 | 2 |
| Vegfa | 1.49E-16 | 0.30662449 | 0.293 | 0.22 | 1.43E-13 | 2 |
| Clu | 1.53E-16 | 0.36000029 | 0.231 | 0.166 | 1.47E-13 | 2 |
| Mef2a | 1.58E-16 | 0.38315029 | 0.157 | 0.103 | 1.52E-13 | 2 |
| App | 3.69E-16 | 0.31998031 | 0.238 | 0.173 | 3.53E-13 | 2 |
| Pfkl | 4.61E-16 | 0.32317129 | 0.217 | 0.154 | 4.42E-13 | 2 |
| Mid1 | 5.46E-16 | 0.30292744 | 0.248 | 0.181 | 5.24E-13 | 2 |
| Flt1 | 7.84E-16 | 0.38799335 | 0.112 | 0.067 | 7.52E-13 | 2 |
| Lgr5 | 1.05E-15 | 0.39532739 | 0.135 | 0.086 | 1.01E-12 | 2 |
| Atp6v0a1 | 4.12E-15 | 0.33778433 | 0.134 | 0.085 | 3.95E-12 | 2 |
| Sem1 | 7.47E-15 | -0.3319233 | 0.4 | 0.458 | 7.16E-12 | 2 |
| Smad2 | 7.72E-15 | 0.30111398 | 0.184 | 0.128 | 7.41E-12 | 2 |
| Pcsk6 | 1.06E-14 | 0.34811387 | 0.118 | 0.074 | 1.02E-11 | 2 |
| Wdr59 | 2.31E-14 | 0.31734994 | 0.148 | 0.099 | 2.22E-11 | 2 |
| Sv2a | 2.59E-14 | 0.37537157 | 0.13 | 0.084 | 2.48E-11 | 2 |
| Slc1a3 | 3.47E-14 | 0.31150966 | 0.139 | 0.091 | 3.33E-11 | 2 |
| Sstr4 | 6.10E-14 | 0.35240269 | 0.109 | 0.067 | 5.85E-11 | 2 |
| Stk3 | 3.04E-13 | 0.31825679 | 0.155 | 0.106 | 2.92E-10 | 2 |
| Rtn4 | 3.98E-13 | 0.28223101 | 0.276 | 0.213 | 3.82E-10 | 2 |
| Slc2a1 | 7.18E-13 | 0.27435865 | 0.174 | 0.122 | 6.88E-10 | 2 |
| Mapk14 | 8.67E-13 | 0.29115954 | 0.197 | 0.143 | 8.31E-10 | 2 |
| Trp53 | 1.12E-12 | 0.31806162 | 0.162 | 0.114 | 1.07E-09 | 2 |
| Bmpr2 | 1.83E-12 | 0.26692671 | 0.214 | 0.159 | 1.75E-09 | 2 |
| Ltbp1 | 4.11E-12 | 0.28584188 | 0.173 | 0.123 | 3.94E-09 | 2 |
| Dnm1l | 6.05E-12 | 0.28486324 | 0.206 | 0.154 | 5.80E-09 | 2 |
| Mib1 | 6.74E-12 | 0.26969874 | 0.182 | 0.132 | 6.46E-09 | 2 |
| Ntrk1 | 7.12E-12 | 0.34049591 | 0.135 | 0.093 | 6.83E-09 | 2 |
| Sv2b | 8.88E-12 | 0.3155225 | 0.11 | 0.071 | 8.52E-09 | 2 |
| Cd14 | 1.70E-11 | 0.29096639 | 0.1 | 0.064 | 1.63E-08 | 2 |
| Penk | 1.77E-11 | 0.29281489 | 0.172 | 0.125 | 1.70E-08 | 2 |
| Atr | 1.88E-11 | 0.30629723 | 0.119 | 0.079 | 1.81E-08 | 2 |
| Anapc16 | 2.09E-11 | 0.29598321 | 0.174 | 0.127 | 2.00E-08 | 2 |
| Id2 | 2.29E-11 | 0.2705256 | 0.176 | 0.128 | 2.20E-08 | 2 |
| P2rx7 | 4.99E-11 | 0.32668474 | 0.104 | 0.068 | 4.79E-08 | 2 |
| Btrc | 6.45E-11 | 0.28561371 | 0.126 | 0.086 | 6.18E-08 | 2 |
| Nsg1 | 1.32E-10 | 0.25125839 | 0.26 | 0.206 | 1.27E-07 | 2 |
| Samhd1 | 2.71E-10 | 0.27454497 | 0.171 | 0.127 | 2.60E-07 | 2 |
| Calm1 | 2.77E-10 | -0.275093 | 0.434 | 0.48 | 2.65E-07 | 2 |
| Irs2 | 3.36E-10 | 0.26260053 | 0.142 | 0.101 | 3.23E-07 | 2 |
| Ctnnb1 | 3.66E-10 | -0.3052343 | 0.192 | 0.241 | 3.51E-07 | 2 |
| Epha4 | 3.76E-10 | -0.3227967 | 0.169 | 0.217 | 3.60E-07 | 2 |
| Pten | 4.04E-10 | 0.30621901 | 0.198 | 0.152 | 3.87E-07 | 2 |
| Cck | 4.41E-10 | 0.2770085 | 0.12 | 0.083 | 4.23E-07 | 2 |
| Tnc | 4.55E-10 | 0.26905202 | 0.206 | 0.158 | 4.37E-07 | 2 |
| Pak2 | 4.66E-10 | 0.25978088 | 0.174 | 0.129 | 4.47E-07 | 2 |
| Tnrc6a | 6.30E-10 | 0.25702624 | 0.157 | 0.115 | 6.04E-07 | 2 |
| Hspa1a | 7.75E-10 | 0.27239245 | 0.102 | 0.068 | 7.44E-07 | 2 |
| Myo6 | 8.91E-10 | 0.31089087 | 0.11 | 0.076 | 8.54E-07 | 2 |
| Tjp1 | 1.02E-09 | 0.25260168 | 0.153 | 0.112 | 9.74E-07 | 2 |
| Cacna1b | 1.55E-09 | 0.26671758 | 0.113 | 0.078 | 1.48E-06 | 2 |
| Ube3a | 2.05E-09 | 0.25451744 | 0.178 | 0.135 | 1.96E-06 | 2 |
| Rab1a | 4.10E-09 | -0.2869832 | 0.309 | 0.356 | 3.93E-06 | 2 |
| Lpin1 | 4.86E-09 | 0.26961729 | 0.108 | 0.075 | 4.66E-06 | 2 |
| Gpr37l1 | 1.15E-08 | 0.27748652 | 0.1 | 0.069 | 1.11E-05 | 2 |
| Cd9 | 1.37E-08 | 0.25631909 | 0.121 | 0.087 | 1.32E-05 | 2 |
| Slc11a2 | 1.57E-08 | 0.25362641 | 0.155 | 0.116 | 1.50E-05 | 2 |
| Tgfb2 | 6.66E-08 | 0.25188328 | 0.158 | 0.121 | 6.38E-05 | 2 |
| Apod | 1.27E-07 | 0.25397811 | 0.104 | 0.075 | 0.00012225 | 2 |
| Gja1 | 2.66E-06 | -0.2525777 | 0.105 | 0.135 | 0.00255245 | 2 |
| Acta2 | 6.96E-06 | -0.2780681 | 0.154 | 0.185 | 0.00667192 | 2 |
| Gnas | 5.67E-110 | -0.4514779 | 0.887 | 0.943 | 5.44E-107 | 3 |
| Col1a1 | 3.16E-100 | 0.28465186 | 1 | 0.998 | 3.03E-97 | 3 |
| Ndrg1 | 1.88E-72 | 0.74690326 | 0.338 | 0.18 | 1.80E-69 | 3 |
| Pcsk6 | 4.30E-68 | 0.47872615 | 0.654 | 0.48 | 4.13E-65 | 3 |
| Tuba1a/b/c | 2.19E-61 | 0.38564205 | 0.816 | 0.681 | 2.10E-58 | 3 |
| Aldoa | 5.19E-58 | 0.59902008 | 0.532 | 0.384 | 4.98E-55 | 3 |
| Cfh | 1.92E-38 | 0.42214684 | 0.511 | 0.385 | 1.84E-35 | 3 |
| Dnm1 | 6.02E-35 | 0.46626383 | 0.36 | 0.245 | 5.77E-32 | 3 |
| Vegfa | 6.32E-33 | 0.47922129 | 0.35 | 0.242 | 6.06E-30 | 3 |
| Cd9 | 3.49E-30 | 0.50606072 | 0.215 | 0.128 | 3.34E-27 | 3 |
| Ldha | 8.42E-28 | 0.45704649 | 0.427 | 0.326 | 8.08E-25 | 3 |
| Serinc3 | 8.70E-26 | -0.4123075 | 0.361 | 0.454 | 8.34E-23 | 3 |
| Cnr1 | 1.04E-24 | 0.46046479 | 0.185 | 0.111 | 9.93E-22 | 3 |
| Hspa8 | 1.63E-24 | -0.3875106 | 0.448 | 0.538 | 1.56E-21 | 3 |
| Mef2a | 7.58E-24 | 0.39979705 | 0.306 | 0.216 | 7.27E-21 | 3 |
| Rab1a | 2.71E-23 | -0.3852082 | 0.433 | 0.516 | 2.60E-20 | 3 |
| Pltp | 1.08E-22 | 0.42064032 | 0.199 | 0.126 | 1.03E-19 | 3 |
| Col3a1 | 1.61E-22 | -0.4651199 | 0.297 | 0.385 | 1.54E-19 | 3 |
| Atpif1 | 2.61E-22 | 0.38165398 | 0.317 | 0.23 | 2.50E-19 | 3 |
| Pfkl | 5.45E-22 | 0.37463387 | 0.32 | 0.232 | 5.22E-19 | 3 |
| Bin1 | 2.16E-21 | 0.39806138 | 0.252 | 0.173 | 2.07E-18 | 3 |
| Ndufa4 | 1.56E-20 | -0.3742812 | 0.332 | 0.415 | 1.49E-17 | 3 |
| Smurf1 | 8.22E-20 | 0.42034209 | 0.152 | 0.092 | 7.88E-17 | 3 |
| Slc2a13 | 1.08E-19 | 0.41642451 | 0.154 | 0.094 | 1.03E-16 | 3 |
| Lifr | 7.95E-19 | 0.31599757 | 0.35 | 0.266 | 7.62E-16 | 3 |
| Mid1 | 1.41E-18 | 0.41110269 | 0.202 | 0.135 | 1.36E-15 | 3 |
| Tubb5 | 2.17E-18 | 0.26467327 | 0.615 | 0.531 | 2.08E-15 | 3 |
| Prex1 | 2.35E-18 | -0.3362632 | 0.355 | 0.434 | 2.26E-15 | 3 |
| Cpe | 5.61E-18 | 0.26732264 | 0.578 | 0.494 | 5.38E-15 | 3 |
| C5ar1 | 1.56E-17 | 0.40371642 | 0.124 | 0.073 | 1.49E-14 | 3 |
| Smad2 | 2.18E-17 | 0.39983977 | 0.153 | 0.097 | 2.09E-14 | 3 |
| Bcl2 | 4.19E-17 | 0.37093368 | 0.131 | 0.078 | 4.02E-14 | 3 |
| Akt1 | 5.76E-17 | 0.27810474 | 0.398 | 0.314 | 5.53E-14 | 3 |
| Serpine2 | 9.07E-17 | 0.31493045 | 0.359 | 0.281 | 8.70E-14 | 3 |
| Map1b | 2.37E-16 | 0.38142214 | 0.171 | 0.113 | 2.28E-13 | 3 |
| Gpr37l1 | 2.72E-16 | 0.36852792 | 0.125 | 0.075 | 2.61E-13 | 3 |
| Selenow | 6.90E-16 | 0.32512186 | 0.238 | 0.171 | 6.62E-13 | 3 |
| Fyn | 1.62E-14 | -0.3010372 | 0.471 | 0.531 | 1.55E-11 | 3 |
| Cd55 | 7.94E-14 | 0.31256125 | 0.181 | 0.126 | 7.61E-11 | 3 |
| Hipk2 | 1.65E-13 | 0.27158432 | 0.381 | 0.31 | 1.58E-10 | 3 |
| Ctsd | 1.99E-13 | -0.3532586 | 0.226 | 0.287 | 1.91E-10 | 3 |
| Slc17a7 | 4.01E-13 | 0.32641711 | 0.108 | 0.066 | 3.84E-10 | 3 |
| Nsg1 | 9.53E-13 | 0.30806839 | 0.153 | 0.104 | 9.14E-10 | 3 |
| Gab2 | 1.69E-12 | 0.31480043 | 0.109 | 0.068 | 1.62E-09 | 3 |
| Ctxn1 | 2.73E-12 | 0.33209522 | 0.151 | 0.104 | 2.62E-09 | 3 |
| Cd6 | 6.14E-12 | 0.31197266 | 0.103 | 0.065 | 5.89E-09 | 3 |
| Slc12a2 | 7.34E-12 | 0.30222069 | 0.164 | 0.115 | 7.04E-09 | 3 |
| Drd1 | 1.79E-11 | 0.30715064 | 0.1 | 0.063 | 1.71E-08 | 3 |
| Atp2b1 | 1.97E-11 | -0.2870465 | 0.436 | 0.486 | 1.89E-08 | 3 |
| Hspa1a | 2.40E-11 | 0.30834863 | 0.104 | 0.066 | 2.30E-08 | 3 |
| Cacna1b | 3.51E-11 | 0.29114468 | 0.182 | 0.133 | 3.36E-08 | 3 |
| Trp53 | 6.55E-11 | 0.30104605 | 0.137 | 0.095 | 6.28E-08 | 3 |
| P2rx7 | 7.66E-11 | 0.3025718 | 0.12 | 0.08 | 7.35E-08 | 3 |
| Slc2a1 | 9.44E-11 | 0.26532641 | 0.169 | 0.122 | 9.05E-08 | 3 |
| Cck | 1.16E-10 | 0.30763474 | 0.14 | 0.098 | 1.11E-07 | 3 |
| Sod1 | 1.23E-10 | -0.2584912 | 0.473 | 0.519 | 1.18E-07 | 3 |
| Wac | 1.67E-10 | 0.25151812 | 0.257 | 0.202 | 1.60E-07 | 3 |
| Aqp4 | 2.79E-10 | 0.29295695 | 0.111 | 0.074 | 2.67E-07 | 3 |
| Pik3cb | 3.14E-10 | 0.2932634 | 0.117 | 0.079 | 3.02E-07 | 3 |
| Trim2 | 3.69E-10 | 0.27328363 | 0.15 | 0.107 | 3.54E-07 | 3 |
| Slc11a2 | 1.26E-09 | 0.27223012 | 0.168 | 0.125 | 1.21E-06 | 3 |
| Epb41l3 | 1.60E-09 | 0.25095466 | 0.248 | 0.197 | 1.53E-06 | 3 |
| Pbx1 | 1.95E-09 | -0.2956023 | 0.228 | 0.278 | 1.87E-06 | 3 |
| Creb1 | 2.15E-09 | 0.25077528 | 0.213 | 0.165 | 2.06E-06 | 3 |
| Cartpt | 2.63E-09 | 0.27667093 | 0.101 | 0.067 | 2.52E-06 | 3 |
| Psen1 | 4.29E-09 | 0.26069827 | 0.173 | 0.13 | 4.12E-06 | 3 |
| Sv2a | 4.90E-09 | 0.27370876 | 0.131 | 0.093 | 4.70E-06 | 3 |
| Mobp | 5.77E-09 | 0.250853 | 0.105 | 0.071 | 5.54E-06 | 3 |
| Idnk | 6.46E-09 | 0.28234427 | 0.114 | 0.08 | 6.20E-06 | 3 |
| Traf3 | 3.37E-08 | 0.26434102 | 0.108 | 0.075 | 3.24E-05 | 3 |
| Gja1 | 3.49E-08 | -0.2525692 | 0.176 | 0.22 | 3.35E-05 | 3 |
| Tnrc6b | 5.04E-08 | -0.266664 | 0.182 | 0.226 | 4.83E-05 | 3 |
| Wdr59 | 8.53E-08 | 0.25202347 | 0.128 | 0.094 | 8.18E-05 | 3 |
| Robo1 | 2.19E-07 | -0.2508989 | 0.185 | 0.227 | 0.00021049 | 3 |
| Cd109 | 4.06E-07 | 0.25352402 | 0.229 | 0.187 | 0.00038918 | 3 |
| Psme4 | 5.31E-07 | -0.2634561 | 0.23 | 0.271 | 0.00050951 | 3 |
| Itm2a | 1.89E-06 | -0.2537229 | 0.146 | 0.182 | 0.00181322 | 3 |
| Gnas | 2.09E-57 | -0.5057817 | 0.909 | 0.961 | 2.01E-54 | 4 |
| Flt1 | 3.15E-37 | 0.66687362 | 0.617 | 0.459 | 3.02E-34 | 4 |
| Aplnr | 6.73E-23 | -0.6518484 | 0.366 | 0.487 | 6.45E-20 | 4 |
| Calm1 | 4.06E-22 | -0.4195464 | 0.66 | 0.744 | 3.89E-19 | 4 |
| Emcn | 1.06E-17 | 0.42792705 | 0.513 | 0.387 | 1.01E-14 | 4 |
| Sparc | 1.79E-17 | -0.3269699 | 0.749 | 0.806 | 1.72E-14 | 4 |
| Aldoa | 1.05E-15 | 0.3944879 | 0.525 | 0.401 | 1.01E-12 | 4 |
| Hspa8 | 5.79E-14 | -0.3620157 | 0.594 | 0.671 | 5.55E-11 | 4 |
| Bin1 | 8.69E-14 | 0.49106907 | 0.29 | 0.192 | 8.33E-11 | 4 |
| Mid1 | 1.32E-11 | 0.4222664 | 0.24 | 0.157 | 1.26E-08 | 4 |
| Vegfa | 3.58E-11 | 0.47087707 | 0.131 | 0.07 | 3.43E-08 | 4 |
| Smurf1 | 1.64E-10 | 0.44681406 | 0.207 | 0.134 | 1.57E-07 | 4 |
| Bmpr2 | 1.84E-10 | 0.38869347 | 0.311 | 0.226 | 1.77E-07 | 4 |
| P2rx7 | 9.28E-10 | 0.44894842 | 0.129 | 0.073 | 8.90E-07 | 4 |
| Cd55 | 2.00E-09 | 0.39205561 | 0.169 | 0.105 | 1.92E-06 | 4 |
| Pde8a | 2.72E-09 | 0.37567422 | 0.293 | 0.216 | 2.61E-06 | 4 |
| Wdr59 | 3.04E-09 | 0.42969126 | 0.184 | 0.119 | 2.91E-06 | 4 |
| Smad2 | 3.19E-09 | 0.34044462 | 0.205 | 0.135 | 3.06E-06 | 4 |
| Grb10 | 4.12E-09 | -0.3289438 | 0.507 | 0.57 | 3.95E-06 | 4 |
| Phc2 | 7.59E-09 | 0.32576321 | 0.317 | 0.236 | 7.28E-06 | 4 |
| Rgs6 | 1.56E-08 | 0.38686166 | 0.101 | 0.055 | 1.49E-05 | 4 |
| Calm2 | 2.25E-08 | -0.2585174 | 0.653 | 0.697 | 2.16E-05 | 4 |
| Mib1 | 3.47E-08 | 0.35540047 | 0.238 | 0.169 | 3.33E-05 | 4 |
| Aplp2 | 9.68E-08 | 0.28741848 | 0.486 | 0.406 | 9.29E-05 | 4 |
| Serpine2 | 1.14E-07 | 0.47297699 | 0.171 | 0.116 | 0.00010902 | 4 |
| Id1 | 1.22E-07 | 0.33754785 | 0.3 | 0.229 | 0.00011684 | 4 |
| Atxn1 | 1.72E-07 | 0.35847944 | 0.115 | 0.069 | 0.00016535 | 4 |
| Tjp1 | 2.62E-07 | 0.26256449 | 0.426 | 0.35 | 0.00025123 | 4 |
| C5ar1 | 3.96E-07 | 0.35952789 | 0.107 | 0.064 | 0.00038016 | 4 |
| Slc17a7 | 8.73E-07 | 0.31631941 | 0.106 | 0.064 | 0.00083729 | 4 |
| Pcsk6 | 9.55E-07 | 0.34945587 | 0.132 | 0.086 | 0.00091578 | 4 |
| Sptbn1 | 1.00E-06 | -0.2941484 | 0.401 | 0.467 | 0.00096012 | 4 |
| Lpar2 | 1.16E-06 | 0.35810684 | 0.112 | 0.07 | 0.00111506 | 4 |
| Btrc | 1.20E-06 | 0.36756448 | 0.133 | 0.087 | 0.00115008 | 4 |
| Ndufa4 | 1.38E-06 | -0.2656035 | 0.288 | 0.358 | 0.00132073 | 4 |
| Pak2 | 1.95E-06 | 0.27507932 | 0.241 | 0.18 | 0.00186715 | 4 |
| Wac | 2.20E-06 | 0.29871641 | 0.267 | 0.205 | 0.00210932 | 4 |
| Hipk2 | 2.29E-06 | 0.37857742 | 0.165 | 0.115 | 0.00219162 | 4 |
| Tgfb1 | 3.46E-06 | 0.27039592 | 0.25 | 0.19 | 0.00332087 | 4 |
| Pten | 3.77E-06 | 0.3390289 | 0.239 | 0.183 | 0.00361183 | 4 |
| Cd9 | 5.76E-06 | 0.3105889 | 0.223 | 0.169 | 0.00551928 | 4 |
| Atpif1 | 6.84E-06 | 0.26315058 | 0.354 | 0.29 | 0.00656427 | 4 |
| Ntrk1 | 7.03E-06 | 0.26884926 | 0.122 | 0.08 | 0.0067396 | 4 |
| Skil | 1.10E-05 | 0.28376547 | 0.238 | 0.184 | 0.01058335 | 4 |
| Slc2a13 | 1.21E-05 | 0.29731242 | 0.1 | 0.063 | 0.01155937 | 4 |
| Apoe | 1.23E-05 | 0.29245853 | 0.179 | 0.13 | 0.01177972 | 4 |
| Sbf2 | 1.24E-05 | 0.25286087 | 0.207 | 0.155 | 0.01189273 | 4 |
| Slco3a1 | 1.24E-05 | 0.31559384 | 0.134 | 0.091 | 0.01192952 | 4 |
| Mef2a | 1.31E-05 | 0.28886155 | 0.222 | 0.169 | 0.0125721 | 4 |
| Stk3 | 1.43E-05 | 0.31452445 | 0.155 | 0.11 | 0.01368501 | 4 |
| Sptan1 | 1.47E-05 | -0.2892735 | 0.317 | 0.375 | 0.01413334 | 4 |
| Efna5 | 1.56E-05 | 0.28708835 | 0.169 | 0.122 | 0.01497964 | 4 |
| Lilrb4a/b | 1.96E-05 | 0.31939866 | 0.137 | 0.095 | 0.01875764 | 4 |
| Ctxn1 | 2.14E-05 | 0.28107956 | 0.231 | 0.178 | 0.02055554 | 4 |
| Jak2 | 2.25E-05 | 0.26717989 | 0.198 | 0.147 | 0.02153496 | 4 |
| Mertk | 2.59E-05 | 0.28849727 | 0.123 | 0.084 | 0.0248844 | 4 |
| Psen1 | 2.89E-05 | 0.3012935 | 0.138 | 0.097 | 0.02775613 | 4 |
| Tnrc6a | 3.94E-05 | 0.28006974 | 0.161 | 0.118 | 0.03782101 | 4 |
| Lrp6 | 3.97E-05 | 0.30828147 | 0.165 | 0.121 | 0.03810979 | 4 |
| Pfkl | 4.16E-05 | 0.29857821 | 0.215 | 0.167 | 0.03993503 | 4 |
| Ngfr | 4.27E-05 | 0.30679362 | 0.13 | 0.091 | 0.04099439 | 4 |
| Hspa8 | 2.51E-40 | -0.619068 | 0.581 | 0.735 | 2.40E-37 | 5 |
| Aldoa | 2.64E-29 | 0.58838523 | 0.659 | 0.519 | 2.54E-26 | 5 |
| Gnas | 2.15E-18 | -0.4059531 | 0.639 | 0.73 | 2.07E-15 | 5 |
| Ctnnb1 | 1.56E-16 | -0.4537269 | 0.416 | 0.537 | 1.50E-13 | 5 |
| Tmsb10 | 2.43E-15 | -0.2558194 | 0.927 | 0.95 | 2.33E-12 | 5 |
| Slc2a1 | 4.79E-15 | 0.46547346 | 0.499 | 0.384 | 4.60E-12 | 5 |
| Ndrg1 | 5.49E-14 | 0.59762613 | 0.249 | 0.15 | 5.27E-11 | 5 |
| Sox9 | 2.57E-10 | -0.5429095 | 0.15 | 0.224 | 2.46E-07 | 5 |
| Ldha | 3.40E-09 | 0.33133294 | 0.586 | 0.512 | 3.26E-06 | 5 |
| Fth1 | 6.06E-09 | -0.2663478 | 0.536 | 0.618 | 5.82E-06 | 5 |
| Vegfa | 1.15E-08 | 0.40107144 | 0.311 | 0.23 | 1.10E-05 | 5 |
| Ywhag | 6.76E-07 | -0.302471 | 0.298 | 0.369 | 0.00064815 | 5 |
| Fgfr2 | 1.98E-06 | -0.3912781 | 0.166 | 0.224 | 0.00189529 | 5 |
| Bsg | 2.79E-06 | 0.28440822 | 0.409 | 0.339 | 0.00267394 | 5 |
| Lpin1 | 3.41E-06 | 0.38590687 | 0.142 | 0.093 | 0.0032714 | 5 |
| Pfkl | 3.66E-06 | 0.31633025 | 0.362 | 0.297 | 0.00350675 | 5 |
| Mid1 | 4.34E-06 | 0.39691142 | 0.26 | 0.2 | 0.00416365 | 5 |
| Map1b | 4.47E-06 | 0.36635673 | 0.264 | 0.204 | 0.00428356 | 5 |
| Usp15 | 5.08E-06 | 0.36214598 | 0.222 | 0.164 | 0.00487579 | 5 |
| Bcl2 | 9.40E-06 | 0.32598044 | 0.177 | 0.124 | 0.0090122 | 5 |
| Spock1 | 2.05E-05 | 0.38602851 | 0.114 | 0.072 | 0.0196772 | 5 |
| Crip1 | 2.74E-05 | 0.36089292 | 0.193 | 0.143 | 0.02622936 | 5 |
| Ghr | 4.03E-05 | 0.27712695 | 0.134 | 0.09 | 0.03862649 | 5 |
| Atr | 5.16E-05 | 0.28751447 | 0.166 | 0.118 | 0.04946003 | 5 |
| Gnas | 5.84E-61 | -0.6547649 | 0.732 | 0.867 | 5.60E-58 | 6 |
| Ptn | 7.65E-48 | -0.7786117 | 0.567 | 0.696 | 7.33E-45 | 6 |
| Col3a1 | 5.58E-42 | -0.5880691 | 0.624 | 0.763 | 5.35E-39 | 6 |
| Col1a1 | 2.48E-33 | -0.417025 | 0.795 | 0.908 | 2.38E-30 | 6 |
| Itm2a | 1.99E-29 | -0.7062897 | 0.346 | 0.502 | 1.91E-26 | 6 |
| Pak1 | 7.89E-27 | 0.8928603 | 0.265 | 0.133 | 7.57E-24 | 6 |
| Col1a2 | 1.61E-26 | -0.4039749 | 0.674 | 0.804 | 1.55E-23 | 6 |
| Egr1 | 4.84E-26 | 0.74946563 | 0.216 | 0.093 | 4.64E-23 | 6 |
| Ngfr | 2.90E-23 | 0.73439843 | 0.233 | 0.113 | 2.78E-20 | 6 |
| Spock1 | 7.51E-23 | 0.737748 | 0.225 | 0.108 | 7.20E-20 | 6 |
| Grb10 | 1.23E-19 | -0.4917197 | 0.484 | 0.602 | 1.18E-16 | 6 |
| Vegfa | 4.07E-18 | 0.54926771 | 0.316 | 0.194 | 3.90E-15 | 6 |
| Sparc | 9.70E-17 | -0.4286954 | 0.38 | 0.507 | 9.30E-14 | 6 |
| Olfm1 | 5.11E-16 | 0.58072775 | 0.204 | 0.11 | 4.90E-13 | 6 |
| Ctbp2 | 9.66E-15 | 0.48290428 | 0.3 | 0.193 | 9.27E-12 | 6 |
| Id1 | 4.68E-14 | 0.53003258 | 0.262 | 0.165 | 4.48E-11 | 6 |
| Skil | 1.01E-13 | 0.50039777 | 0.285 | 0.184 | 9.67E-11 | 6 |
| Tgfb1 | 1.34E-13 | 0.63541159 | 0.162 | 0.085 | 1.29E-10 | 6 |
| Gria2 | 1.55E-13 | 0.67225032 | 0.161 | 0.085 | 1.49E-10 | 6 |
| Mmp2 | 2.96E-13 | -0.5485627 | 0.293 | 0.384 | 2.84E-10 | 6 |
| Serpine2 | 9.51E-13 | 0.49151434 | 0.281 | 0.184 | 9.12E-10 | 6 |
| Igf1r | 2.01E-12 | 0.45635073 | 0.275 | 0.181 | 1.93E-09 | 6 |
| Maged1 | 3.50E-12 | -0.3361935 | 0.603 | 0.672 | 3.36E-09 | 6 |
| Tle4 | 6.28E-12 | 0.5191709 | 0.115 | 0.054 | 6.02E-09 | 6 |
| Gas2 | 1.03E-11 | -0.5960352 | 0.097 | 0.167 | 9.83E-09 | 6 |
| Mid1 | 1.38E-11 | 0.47022488 | 0.249 | 0.163 | 1.33E-08 | 6 |
| Map4k4 | 3.15E-11 | 0.38435565 | 0.357 | 0.259 | 3.02E-08 | 6 |
| Aldoa | 1.81E-10 | 0.37153098 | 0.451 | 0.355 | 1.74E-07 | 6 |
| Mfge8 | 2.47E-10 | 0.40937289 | 0.177 | 0.105 | 2.36E-07 | 6 |
| Srgn | 3.03E-10 | 0.43270614 | 0.115 | 0.058 | 2.91E-07 | 6 |
| Thy1 | 3.34E-10 | 0.40120831 | 0.214 | 0.137 | 3.20E-07 | 6 |
| Ifngr2 | 6.51E-10 | 0.40987588 | 0.223 | 0.145 | 6.24E-07 | 6 |
| Cd9 | 1.38E-09 | 0.42381477 | 0.185 | 0.116 | 1.32E-06 | 6 |
| P2rx7 | 1.70E-09 | 0.3990726 | 0.153 | 0.089 | 1.63E-06 | 6 |
| Slc6a1 | 1.79E-09 | 0.48227647 | 0.102 | 0.052 | 1.72E-06 | 6 |
| Slc7a5 | 1.97E-09 | 0.37373114 | 0.199 | 0.127 | 1.89E-06 | 6 |
| Adgrl2 | 2.21E-09 | 0.38629835 | 0.224 | 0.149 | 2.12E-06 | 6 |
| Pde8a | 2.29E-09 | 0.40221737 | 0.102 | 0.05 | 2.19E-06 | 6 |
| Bmpr2 | 2.30E-09 | 0.31865228 | 0.283 | 0.199 | 2.21E-06 | 6 |
| Pdgfrb | 2.48E-09 | 0.38787672 | 0.218 | 0.144 | 2.38E-06 | 6 |
| Rasa1 | 3.02E-09 | 0.43129286 | 0.286 | 0.207 | 2.89E-06 | 6 |
| Ptpn1 | 3.62E-09 | 0.4211563 | 0.217 | 0.145 | 3.47E-06 | 6 |
| Gnai2 | 7.02E-09 | 0.28611043 | 0.458 | 0.361 | 6.73E-06 | 6 |
| Fbxw7 | 2.40E-08 | 0.36328247 | 0.12 | 0.067 | 2.30E-05 | 6 |
| Sbf2 | 5.42E-08 | 0.39137507 | 0.179 | 0.117 | 5.20E-05 | 6 |
| Acer3 | 7.22E-08 | 0.38392443 | 0.11 | 0.061 | 6.92E-05 | 6 |
| Ncor2 | 7.41E-08 | 0.3349 | 0.278 | 0.204 | 7.11E-05 | 6 |
| Ahcyl1 | 7.86E-08 | 0.26948098 | 0.32 | 0.24 | 7.54E-05 | 6 |
| Slc17a7 | 1.01E-07 | 0.38612535 | 0.105 | 0.058 | 9.67E-05 | 6 |
| Car8 | 1.19E-07 | 0.35571631 | 0.112 | 0.064 | 0.00011372 | 6 |
| Chchd10 | 1.53E-07 | 0.3641127 | 0.154 | 0.098 | 0.00014688 | 6 |
| Aplp2 | 2.18E-07 | 0.33933533 | 0.294 | 0.222 | 0.00020871 | 6 |
| Nlk | 2.53E-07 | 0.39208477 | 0.101 | 0.056 | 0.00024281 | 6 |
| Vim | 2.96E-07 | -0.2608341 | 0.541 | 0.602 | 0.00028421 | 6 |
| P2rx4 | 3.12E-07 | 0.33816479 | 0.131 | 0.08 | 0.0002993 | 6 |
| Trp53 | 4.57E-07 | 0.30716507 | 0.216 | 0.153 | 0.00043868 | 6 |
| Hspa8 | 4.81E-07 | -0.3215138 | 0.497 | 0.543 | 0.00046092 | 6 |
| C1qc | 5.53E-07 | 0.39457057 | 0.111 | 0.066 | 0.00053042 | 6 |
| Epn2 | 5.54E-07 | 0.32164038 | 0.202 | 0.141 | 0.00053083 | 6 |
| Grk3 | 6.00E-07 | 0.36017045 | 0.111 | 0.066 | 0.00057515 | 6 |
| Lpar6 | 6.40E-07 | 0.33347735 | 0.113 | 0.067 | 0.0006139 | 6 |
| Mllt11 | 8.27E-07 | 0.34710808 | 0.165 | 0.111 | 0.00079329 | 6 |
| Cck | 9.45E-07 | 0.34472685 | 0.125 | 0.078 | 0.00090619 | 6 |
| Magi1 | 9.78E-07 | 0.38458006 | 0.138 | 0.089 | 0.00093802 | 6 |
| Hspa1a | 1.06E-06 | 0.34293913 | 0.102 | 0.059 | 0.00101702 | 6 |
| Prkca | 1.28E-06 | 0.3540601 | 0.139 | 0.089 | 0.00122897 | 6 |
| Tcf12 | 1.32E-06 | 0.26809916 | 0.324 | 0.253 | 0.00126374 | 6 |
| Ednra | 1.43E-06 | 0.33718322 | 0.121 | 0.075 | 0.00137558 | 6 |
| Nrgn | 2.12E-06 | 0.34578582 | 0.118 | 0.073 | 0.00203285 | 6 |
| Stk3 | 3.17E-06 | 0.31134239 | 0.166 | 0.113 | 0.00303938 | 6 |
| Gdnf | 3.43E-06 | 0.33076817 | 0.107 | 0.065 | 0.00328729 | 6 |
| Ranbp2 | 3.45E-06 | 0.27770268 | 0.277 | 0.213 | 0.00331263 | 6 |
| Gls | 4.29E-06 | 0.25434958 | 0.237 | 0.175 | 0.00411777 | 6 |
| Prkg1 | 4.34E-06 | 0.29120753 | 0.181 | 0.127 | 0.00416376 | 6 |
| Ambra1 | 4.38E-06 | 0.25172279 | 0.188 | 0.132 | 0.00420462 | 6 |
| Gsn | 4.43E-06 | -0.4356466 | 0.207 | 0.26 | 0.00424678 | 6 |
| Ndrg1 | 4.54E-06 | 0.33671773 | 0.127 | 0.082 | 0.00435097 | 6 |
| Sstr4 | 5.03E-06 | 0.32379717 | 0.128 | 0.083 | 0.00481984 | 6 |
| Hdac9 | 5.16E-06 | 0.28119576 | 0.1 | 0.06 | 0.00494947 | 6 |
| Igfbp7 | 5.29E-06 | 0.30416579 | 0.112 | 0.07 | 0.00506943 | 6 |
| Pten | 5.80E-06 | 0.28156057 | 0.231 | 0.171 | 0.00556305 | 6 |
| Slc11a2 | 5.92E-06 | 0.25574604 | 0.191 | 0.136 | 0.00567282 | 6 |
| Tspan5 | 6.17E-06 | 0.27489935 | 0.109 | 0.067 | 0.00591899 | 6 |
| Sem1 | 6.21E-06 | -0.2947004 | 0.472 | 0.517 | 0.00595342 | 6 |
| Wac | 6.58E-06 | 0.25789036 | 0.297 | 0.232 | 0.00631463 | 6 |
| Cd55 | 7.67E-06 | 0.26630995 | 0.136 | 0.091 | 0.00735353 | 6 |
| Vipr1 | 9.65E-06 | 0.27262523 | 0.112 | 0.071 | 0.00925576 | 6 |
| Itgav | 1.03E-05 | 0.27613786 | 0.145 | 0.098 | 0.00988904 | 6 |
| Fos | 1.04E-05 | 0.29876745 | 0.123 | 0.08 | 0.0099471 | 6 |
| Fzd3 | 1.23E-05 | 0.30420313 | 0.146 | 0.1 | 0.01178172 | 6 |
| Cxcl14 | 1.24E-05 | 0.3922563 | 0.112 | 0.073 | 0.01187922 | 6 |
| Pfkl | 1.24E-05 | 0.29484698 | 0.195 | 0.143 | 0.01189906 | 6 |
| Homer1 | 1.33E-05 | 0.26649691 | 0.178 | 0.127 | 0.01276053 | 6 |
| Pcp2 | 1.50E-05 | 0.31691866 | 0.117 | 0.076 | 0.01436397 | 6 |
| Cblb | 1.69E-05 | 0.25303277 | 0.217 | 0.162 | 0.01622634 | 6 |
| Mt1 | 2.06E-05 | 0.28280457 | 0.107 | 0.068 | 0.01978871 | 6 |
| Wwp2 | 2.08E-05 | -0.3577471 | 0.206 | 0.258 | 0.01992611 | 6 |
| Ube2g1 | 2.78E-05 | 0.27250321 | 0.228 | 0.174 | 0.0266935 | 6 |
| Atg13 | 3.07E-05 | 0.28178053 | 0.128 | 0.087 | 0.02939435 | 6 |
| Dab1 | 3.19E-05 | 0.30865394 | 0.128 | 0.087 | 0.03054537 | 6 |
| Snap25 | 3.56E-05 | 0.33575658 | 0.12 | 0.081 | 0.03412323 | 6 |
| Pde3b | 3.62E-05 | 0.31984446 | 0.13 | 0.089 | 0.03467168 | 6 |
| St3gal6 | 4.31E-05 | 0.29097751 | 0.122 | 0.082 | 0.04130205 | 6 |
| Birc6 | 5.09E-05 | 0.26797112 | 0.235 | 0.182 | 0.04885956 | 6 |
| Gnas | 4.25E-20 | -0.3640619 | 0.844 | 0.904 | 4.08E-17 | 7 |
| Clu | 6.61E-15 | 0.59845176 | 0.337 | 0.217 | 6.34E-12 | 7 |
| Aldoa | 1.75E-13 | 0.46062144 | 0.573 | 0.465 | 1.68E-10 | 7 |
| Hnrnpa2b1 | 1.05E-12 | -0.2798413 | 0.929 | 0.953 | 1.00E-09 | 7 |
| Axl | 2.95E-12 | 0.61947611 | 0.201 | 0.114 | 2.83E-09 | 7 |
| Sox9 | 1.14E-11 | -0.3267339 | 0.539 | 0.643 | 1.09E-08 | 7 |
| Col3a1 | 4.39E-11 | -0.5409803 | 0.449 | 0.52 | 4.21E-08 | 7 |
| Grb10 | 4.47E-10 | -0.3812733 | 0.54 | 0.616 | 4.29E-07 | 7 |
| Hspa8 | 1.35E-09 | -0.4218201 | 0.412 | 0.488 | 1.29E-06 | 7 |
| Avpr1a | 7.32E-09 | 0.48879796 | 0.1 | 0.049 | 7.02E-06 | 7 |
| Hipk2 | 3.52E-08 | 0.49907042 | 0.147 | 0.087 | 3.38E-05 | 7 |
| Bsg | 3.74E-08 | 0.37440939 | 0.387 | 0.299 | 3.59E-05 | 7 |
| Col1a2 | 5.57E-08 | -0.4169878 | 0.596 | 0.629 | 5.34E-05 | 7 |
| Wdr59 | 7.39E-08 | 0.43192637 | 0.163 | 0.1 | 7.09E-05 | 7 |
| Creb5 | 1.43E-07 | 0.47321291 | 0.147 | 0.089 | 0.00013699 | 7 |
| Ctnnb1 | 2.66E-07 | -0.5036381 | 0.163 | 0.231 | 0.00025514 | 7 |
| Sparc | 3.49E-07 | -0.3262138 | 0.563 | 0.624 | 0.00033447 | 7 |
| Ctxn1 | 3.63E-07 | 0.42198878 | 0.183 | 0.12 | 0.00034832 | 7 |
| Rims1 | 3.74E-07 | 0.41259726 | 0.105 | 0.058 | 0.00035852 | 7 |
| Sem1 | 4.34E-07 | -0.4204895 | 0.393 | 0.457 | 0.00041583 | 7 |
| Sstr2 | 5.28E-07 | 0.45070951 | 0.115 | 0.066 | 0.00050588 | 7 |
| Ptn | 9.17E-07 | -0.507371 | 0.253 | 0.32 | 0.00087923 | 7 |
| Bin1 | 9.51E-07 | 0.48205948 | 0.185 | 0.126 | 0.00091211 | 7 |
| Cck | 1.00E-06 | 0.49343758 | 0.139 | 0.087 | 0.00096255 | 7 |
| Rora | 1.43E-06 | 0.36928073 | 0.155 | 0.1 | 0.00137013 | 7 |
| Sv2a | 1.55E-06 | 0.39777917 | 0.118 | 0.07 | 0.00148295 | 7 |
| Mmp2 | 1.77E-06 | -0.4108366 | 0.222 | 0.292 | 0.00169875 | 7 |
| Sorbs1 | 1.77E-06 | 0.39502944 | 0.278 | 0.211 | 0.00170008 | 7 |
| P2rx7 | 2.73E-06 | 0.42584025 | 0.124 | 0.076 | 0.00261364 | 7 |
| Mib1 | 4.89E-06 | 0.33203489 | 0.166 | 0.111 | 0.00468974 | 7 |
| Tcf12 | 6.08E-06 | 0.33881312 | 0.26 | 0.195 | 0.005827 | 7 |
| Atpif1 | 6.50E-06 | 0.27695438 | 0.265 | 0.197 | 0.00623576 | 7 |
| Dync1h1 | 1.43E-05 | 0.39676348 | 0.21 | 0.155 | 0.01374754 | 7 |
| Mertk | 1.61E-05 | 0.36669496 | 0.18 | 0.128 | 0.01547022 | 7 |
| Bcl2 | 1.81E-05 | 0.32652472 | 0.136 | 0.089 | 0.01733858 | 7 |
| Tnrc6a | 1.93E-05 | 0.35621822 | 0.14 | 0.093 | 0.01853736 | 7 |
| Zbtb20 | 2.55E-05 | 0.37341643 | 0.138 | 0.093 | 0.02441156 | 7 |
| Pak2 | 3.63E-05 | 0.31216632 | 0.152 | 0.104 | 0.03484435 | 7 |
| Nsg1 | 3.85E-05 | 0.33203865 | 0.181 | 0.13 | 0.03687954 | 7 |
| Lilrb4a/b | 4.72E-05 | 0.37749743 | 0.157 | 0.11 | 0.04526839 | 7 |
| Pten | 4.88E-05 | 0.36952136 | 0.15 | 0.104 | 0.04676226 | 7 |
| Gnas | 6.11E-38 | -0.6203657 | 0.752 | 0.858 | 5.86E-35 | 8 |
| Gria2 | 1.00E-32 | 1.2230932 | 0.243 | 0.089 | 9.59E-30 | 8 |
| Aldoa | 1.43E-25 | 0.68490431 | 0.561 | 0.373 | 1.37E-22 | 8 |
| Col1a1 | 1.85E-20 | 0.38340107 | 0.935 | 0.878 | 1.78E-17 | 8 |
| Adgrl2 | 2.92E-20 | 0.79288695 | 0.309 | 0.17 | 2.80E-17 | 8 |
| Col3a1 | 1.43E-15 | -0.4181458 | 0.748 | 0.815 | 1.37E-12 | 8 |
| Tmsb10 | 2.01E-15 | -0.3147841 | 0.923 | 0.963 | 1.93E-12 | 8 |
| Hspa8 | 5.84E-13 | -0.3892723 | 0.533 | 0.632 | 5.60E-10 | 8 |
| Fus | 2.29E-12 | 0.33380515 | 0.735 | 0.604 | 2.19E-09 | 8 |
| Lifr | 9.95E-11 | 0.55297555 | 0.182 | 0.101 | 9.54E-08 | 8 |
| Nsg1 | 1.83E-10 | 0.451786 | 0.292 | 0.19 | 1.76E-07 | 8 |
| Ldha | 3.21E-10 | 0.44723641 | 0.449 | 0.347 | 3.08E-07 | 8 |
| Slc2a13 | 5.95E-10 | 0.49589933 | 0.103 | 0.046 | 5.70E-07 | 8 |
| Itm2a | 1.55E-09 | -0.4666203 | 0.395 | 0.488 | 1.49E-06 | 8 |
| Wdr59 | 1.47E-08 | 0.47908731 | 0.155 | 0.088 | 1.41E-05 | 8 |
| Atr | 3.46E-08 | 0.45036746 | 0.125 | 0.066 | 3.32E-05 | 8 |
| Cox4i1 | 3.81E-08 | 0.33582578 | 0.436 | 0.331 | 3.66E-05 | 8 |
| Bsg | 4.88E-08 | 0.37570962 | 0.324 | 0.232 | 4.68E-05 | 8 |
| Gnai2 | 7.65E-08 | 0.38615087 | 0.35 | 0.261 | 7.34E-05 | 8 |
| Efna5 | 9.74E-08 | 0.3782354 | 0.253 | 0.171 | 9.35E-05 | 8 |
| P2rx7 | 1.39E-07 | 0.44622451 | 0.113 | 0.06 | 0.00013364 | 8 |
| Cck | 1.86E-07 | 0.43839815 | 0.126 | 0.07 | 0.00017853 | 8 |
| Creb1 | 2.24E-07 | 0.38745882 | 0.223 | 0.149 | 0.00021487 | 8 |
| Grb10 | 2.33E-07 | -0.3234217 | 0.647 | 0.683 | 0.00022375 | 8 |
| Dcn | 2.35E-07 | 0.35546113 | 0.384 | 0.293 | 0.00022546 | 8 |
| Ifngr2 | 2.57E-07 | 0.42241562 | 0.188 | 0.12 | 0.00024655 | 8 |
| Stk3 | 2.66E-07 | 0.4077695 | 0.133 | 0.076 | 0.00025555 | 8 |
| Sv2a | 3.66E-07 | 0.40227455 | 0.147 | 0.088 | 0.00035117 | 8 |
| Mid1 | 3.75E-07 | 0.41059351 | 0.249 | 0.172 | 0.0003595 | 8 |
| Epb41l3 | 4.18E-07 | 0.35045597 | 0.344 | 0.258 | 0.00040131 | 8 |
| Vegfa | 4.65E-07 | 0.3822239 | 0.268 | 0.191 | 0.00044582 | 8 |
| Dnm1 | 5.40E-07 | 0.38397386 | 0.338 | 0.254 | 0.00051803 | 8 |
| Wac | 7.25E-07 | 0.29305214 | 0.3 | 0.215 | 0.00069567 | 8 |
| C5ar1 | 1.03E-06 | 0.40162832 | 0.108 | 0.059 | 0.00099073 | 8 |
| Rbpj | 1.46E-06 | 0.33783014 | 0.406 | 0.325 | 0.00140332 | 8 |
| Slc2a1 | 1.58E-06 | 0.41489026 | 0.17 | 0.109 | 0.00151456 | 8 |
| Pten | 1.96E-06 | 0.39474626 | 0.224 | 0.156 | 0.00187814 | 8 |
| Acta2 | 2.02E-06 | -0.5745524 | 0.183 | 0.252 | 0.00193495 | 8 |
| Anapc16 | 2.88E-06 | 0.36580595 | 0.179 | 0.118 | 0.00276141 | 8 |
| Mapk14 | 2.95E-06 | 0.40928525 | 0.192 | 0.13 | 0.00282545 | 8 |
| Id1 | 3.30E-06 | 0.3878316 | 0.211 | 0.147 | 0.00316658 | 8 |
| Lep | 4.04E-06 | 0.38469129 | 0.121 | 0.072 | 0.00387064 | 8 |
| Dner | 4.25E-06 | 0.34970898 | 0.135 | 0.082 | 0.00407262 | 8 |
| Lrp6 | 4.39E-06 | 0.38782414 | 0.175 | 0.116 | 0.00420896 | 8 |
| Atpif1 | 5.11E-06 | 0.2838604 | 0.348 | 0.266 | 0.00489855 | 8 |
| Sstr4 | 5.45E-06 | 0.36856307 | 0.103 | 0.058 | 0.00522951 | 8 |
| Serpine2 | 5.63E-06 | 0.40128338 | 0.133 | 0.082 | 0.00540063 | 8 |
| Trim2 | 7.91E-06 | 0.37193698 | 0.177 | 0.119 | 0.00758479 | 8 |
| Ndrg1 | 8.23E-06 | 0.37712178 | 0.119 | 0.072 | 0.00788788 | 8 |
| Hipk2 | 8.45E-06 | 0.32398376 | 0.184 | 0.124 | 0.00810426 | 8 |
| Fth1 | 8.97E-06 | -0.3176172 | 0.509 | 0.562 | 0.00860065 | 8 |
| Lpar2 | 9.21E-06 | 0.35235336 | 0.131 | 0.081 | 0.00883122 | 8 |
| Olfm1 | 9.34E-06 | 0.2938007 | 0.146 | 0.093 | 0.00895704 | 8 |
| Slc11a2 | 1.02E-05 | 0.40780331 | 0.164 | 0.11 | 0.00979618 | 8 |
| Grin2a | 1.26E-05 | 0.35820477 | 0.139 | 0.088 | 0.01206683 | 8 |
| Lgr5 | 1.32E-05 | 0.37573324 | 0.211 | 0.15 | 0.01268249 | 8 |
| Gcgr | 1.39E-05 | 0.33347097 | 0.13 | 0.081 | 0.01329799 | 8 |
| Traf3 | 1.39E-05 | 0.36865642 | 0.108 | 0.064 | 0.01337558 | 8 |
| Thy1 | 1.40E-05 | 0.42821606 | 0.128 | 0.08 | 0.01343866 | 8 |
| Bcl2 | 1.45E-05 | 0.39784429 | 0.137 | 0.088 | 0.01390568 | 8 |
| Gpr37l1 | 1.81E-05 | 0.34162606 | 0.105 | 0.062 | 0.01740493 | 8 |
| Tcf12 | 2.15E-05 | 0.28099759 | 0.3 | 0.229 | 0.0206525 | 8 |
| Slc12a2 | 2.79E-05 | 0.30047396 | 0.156 | 0.103 | 0.02678348 | 8 |
| Htt | 4.64E-05 | 0.26478193 | 0.204 | 0.145 | 0.04446023 | 8 |
| Gnas | 2.09E-20 | -0.6467301 | 0.686 | 0.815 | 2.00E-17 | 9 |
| Aldoa | 1.46E-14 | 0.57646419 | 0.638 | 0.476 | 1.40E-11 | 9 |
| Slc11a2 | 1.27E-10 | 0.80697156 | 0.21 | 0.097 | 1.22E-07 | 9 |
| Atp6v0a1 | 3.32E-09 | 0.68260265 | 0.282 | 0.164 | 3.19E-06 | 9 |
| Lilrb4a/b | 6.26E-08 | 0.56925277 | 0.314 | 0.202 | 6.01E-05 | 9 |
| Gnai2 | 1.08E-07 | 0.4421732 | 0.551 | 0.438 | 0.00010363 | 9 |
| Dlk1 | 3.75E-07 | -0.5030536 | 0.258 | 0.367 | 0.00035924 | 9 |
| Col3a1 | 1.13E-06 | -0.3707825 | 0.5 | 0.607 | 0.00108256 | 9 |
| Hipk2 | 2.35E-06 | 0.61271378 | 0.205 | 0.121 | 0.00225165 | 9 |
| Atpif1 | 4.87E-06 | 0.39588529 | 0.432 | 0.334 | 0.00467123 | 9 |
| Serpine2 | 5.01E-06 | 0.52162747 | 0.164 | 0.088 | 0.0048032 | 9 |
| Slc2a1 | 6.72E-06 | 0.60445819 | 0.194 | 0.115 | 0.00644272 | 9 |
| Cd55 | 1.06E-05 | 0.44814633 | 0.149 | 0.079 | 0.01019749 | 9 |
| C3 | 2.22E-05 | 0.57048706 | 0.101 | 0.046 | 0.02125462 | 9 |
| Creb1 | 3.44E-05 | 0.46742738 | 0.218 | 0.14 | 0.03294855 | 9 |
| Gnas | 1.42E-20 | -0.4975806 | 0.837 | 0.904 | 1.36E-17 | 10 |
| Aldoa | 1.09E-13 | 0.66845654 | 0.503 | 0.34 | 1.05E-10 | 10 |
| Tmsb10 | 6.99E-10 | -0.2649969 | 0.937 | 0.974 | 6.71E-07 | 10 |
| Itm2a | 1.39E-09 | -0.6196303 | 0.543 | 0.627 | 1.33E-06 | 10 |
| Bin1 | 2.25E-08 | 0.53377254 | 0.261 | 0.151 | 2.16E-05 | 10 |
| Calm1 | 4.46E-08 | -0.447721 | 0.515 | 0.614 | 4.27E-05 | 10 |
| Dlk1 | 7.96E-08 | 0.54859958 | 0.307 | 0.192 | 7.63E-05 | 10 |
| C5ar1 | 1.18E-07 | 0.55185635 | 0.109 | 0.043 | 0.00011342 | 10 |
| Hspa8 | 1.61E-07 | -0.4132022 | 0.515 | 0.617 | 0.00015463 | 10 |
| Tuba1a/b/c | 2.04E-07 | 0.25871322 | 0.894 | 0.819 | 0.0001958 | 10 |
| Col3a1 | 3.72E-07 | -0.3198076 | 0.747 | 0.817 | 0.0003571 | 10 |
| Cox4i1 | 6.52E-06 | 0.38161205 | 0.422 | 0.321 | 0.00625003 | 10 |
| Grb10 | 6.52E-06 | -0.3357213 | 0.646 | 0.702 | 0.00625677 | 10 |
| Cacna1d | 1.08E-05 | 0.51210717 | 0.133 | 0.07 | 0.01033217 | 10 |
| Adgrl2 | 1.30E-05 | 0.53019207 | 0.158 | 0.09 | 0.01246316 | 10 |
| Pgm1 | 1.34E-05 | 0.49175892 | 0.155 | 0.087 | 0.0128813 | 10 |
| Vcan | 1.36E-05 | 0.37666054 | 0.489 | 0.391 | 0.01299986 | 10 |
| Srgn | 2.33E-05 | 0.4357392 | 0.103 | 0.049 | 0.02234626 | 10 |
| Atpif1 | 2.41E-05 | 0.38149522 | 0.406 | 0.311 | 0.02312922 | 10 |
| Ngfr | 2.77E-05 | 0.48483011 | 0.133 | 0.073 | 0.02656804 | 10 |
| Slc17a7 | 2.96E-05 | 0.48257727 | 0.103 | 0.05 | 0.02839342 | 10 |
| Gnai2 | 3.26E-05 | 0.31683515 | 0.403 | 0.303 | 0.03128201 | 10 |
| Cck | 4.90E-05 | 0.47374177 | 0.147 | 0.085 | 0.04696677 | 10 |
| Col1a1 | 2.85E-20 | -0.6759071 | 0.9 | 0.965 | 2.74E-17 | 11 |
| Gnas | 2.48E-18 | -0.574313 | 0.819 | 0.898 | 2.38E-15 | 11 |
| Col1a2 | 8.44E-18 | -0.6852802 | 0.821 | 0.923 | 8.10E-15 | 11 |
| Sparc | 7.45E-16 | -0.7042359 | 0.672 | 0.823 | 7.14E-13 | 11 |
| Prex1 | 4.91E-11 | -0.9346911 | 0.134 | 0.293 | 4.71E-08 | 11 |
| Ctnnb1 | 3.67E-10 | -0.6794848 | 0.314 | 0.495 | 3.52E-07 | 11 |
| Tubb5 | 1.15E-08 | 0.58538968 | 0.641 | 0.505 | 1.10E-05 | 11 |
| Dapk2 | 5.79E-08 | -0.7261454 | 0.152 | 0.287 | 5.55E-05 | 11 |
| Hmgb1 | 1.91E-07 | 0.41817315 | 0.859 | 0.765 | 0.00018301 | 11 |
| Tuba1a/b/c | 3.75E-07 | 0.40217416 | 0.797 | 0.674 | 0.00035952 | 11 |
| H3f3b | 7.53E-07 | 0.38567119 | 0.829 | 0.786 | 0.00072211 | 11 |
| Cfh | 1.08E-06 | -0.5687882 | 0.309 | 0.445 | 0.00103643 | 11 |
| Cpe | 1.84E-06 | -0.4532919 | 0.338 | 0.484 | 0.00176888 | 11 |
| Thy1 | 1.86E-06 | 0.79457116 | 0.181 | 0.081 | 0.00178181 | 11 |
| Gja1 | 3.46E-06 | -0.6166756 | 0.143 | 0.256 | 0.00331426 | 11 |
| Spp1 | 2.51E-05 | -0.320935 | 1 | 1 | 0.02408788 | 11 |
| Ybx1 | 2.60E-05 | 0.39571667 | 0.621 | 0.495 | 0.02497419 | 11 |
| Mef2c | 3.92E-05 | -0.3974213 | 0.336 | 0.474 | 0.03757023 | 11 |
| Gnas | 1.18E-08 | -0.5217103 | 0.739 | 0.844 | 1.13E-05 | 12 |
| Dbi | 5.86E-08 | -0.5447295 | 0.748 | 0.823 | 5.62E-05 | 12 |
| Plp1 | 2.87E-07 | 0.59513766 | 0.543 | 0.371 | 0.00027492 | 12 |
| Vcan | 1.17E-06 | 0.679178 | 0.57 | 0.423 | 0.00111989 | 12 |
| Tmsb10 | 1.53E-06 | -0.3315119 | 0.932 | 0.977 | 0.00146331 | 12 |
| Aldoa | 1.74E-06 | 0.59135693 | 0.448 | 0.288 | 0.00167197 | 12 |
| Gap43 | 2.46E-06 | 0.6659831 | 0.368 | 0.219 | 0.00236297 | 12 |
| Lrp6 | 2.53E-06 | 0.69109913 | 0.214 | 0.095 | 0.0024223 | 12 |
| Gab2 | 1.21E-05 | 0.68991672 | 0.157 | 0.063 | 0.01160507 | 12 |
| Prnp | 1.62E-05 | 0.67661248 | 0.196 | 0.091 | 0.01555955 | 12 |
| Grb10 | 1.74E-05 | -0.5078384 | 0.439 | 0.581 | 0.01668267 | 12 |
| Hspa8 | 2.22E-05 | -0.5477963 | 0.418 | 0.558 | 0.02127187 | 12 |
| Traf3 | 4.94E-05 | 0.706988 | 0.107 | 0.036 | 0.04735401 | 12 |
| Gnas | 4.52E-08 | -0.5633998 | 0.927 | 0.949 | 4.34E-05 | 13 |
| Flt1 | 8.45E-08 | 1.03961896 | 0.497 | 0.253 | 8.11E-05 | 13 |
| Scg2 | 2.12E-06 | 1.04330249 | 0.162 | 0.03 | 0.00203733 | 13 |
| Tagln | 2.25E-06 | 0.65121284 | 0.659 | 0.392 | 0.00215448 | 13 |
| Vim | 3.05E-06 | 0.50961713 | 0.911 | 0.789 | 0.00292779 | 13 |
| Rgs6 | 9.84E-06 | 0.96714234 | 0.123 | 0.017 | 0.00943974 | 13 |
| Ctxn1 | 1.23E-05 | 0.76918927 | 0.425 | 0.219 | 0.01177302 | 13 |
| Fbn1 | 3.65E-05 | 0.56170443 | 0.536 | 0.316 | 0.03496824 | 13 |
| Rgs5 | 3.85E-05 | -0.4498447 | 0.659 | 0.819 | 0.03692894 | 13 |
| Arhgef7 | 4.34E-05 | 0.68366766 | 0.391 | 0.203 | 0.04161126 | 13 |
| Emcn | 5.10E-05 | 0.89697408 | 0.235 | 0.089 | 0.04889644 | 13 |
| Tank | 3.71E-06 | -1.8795805 | 0.077 | 0.364 | 0.00355869 | 14 |
| Dnm3 | 6.86E-06 | 1.10771117 | 0.169 | 0.032 | 0.00657979 | 14 |
| Vim | 8.35E-06 | -0.8876909 | 0.615 | 0.805 | 0.00800398 | 14 |
| Csf1r | 4.01E-05 | -1.3791982 | 0.2 | 0.466 | 0.03845006 | 14 |
